# Tunable Magnetorheological Elastomer Substrates for Studying Mechano-Genetic Interactions in Hypertrophic Cardiomyopathy

**DOI:** 10.64898/2026.10.24.761361

**Authors:** Yasaman Kargar Gaz Kooh, Yuwen Sun, Ganesh Malayath, Leah Hayem, Bahareh Bahmani, Nongmaithem Debeni Devi, Huanzhu Jiang, Feng Long, Sharon Cresci, Guy M. Genin, Nathaniel Huebsch

## Abstract

Hypertrophic cardiomyopathy (HCM) is the most common inherited cardiac disease. While sarcomeric genomic variants drive HCM, a role for mechanical stress in disease pathogenesis is increasingly appreciated. However, existing *in vitro* models of HCM rely on model systems (e.g., hydrogels) with static mechanical properties, precluding investigation of how dynamic mechanical loading interacts with genetic susceptibility during disease initiation. Here, we leveraged a dynamically tunable magnetorheological elastomer (MRE) platform to model progressive mechanical stiffening in order to study mechano-genetic interactions in human induced pluripotent stem cell (hiPSC) derived cardiomyocytes harboring an HCM-associated MYBPC3 frameshift (*MYBPC3*fs) loss-of-function variant. Dynamic stiffening selectively unmasked pathological structural remodeling in *MYBPC3*fs cardiomyocytes while producing only modest changes in isogenic controls, suggesting that *MYBPC3*fs confers heightened mechanosensitivity. In *MYBPC3*fs cardiomyocytes, mechanical stiffening drove hypertrophic remodeling, sarcomere disorganization, nuclear remodeling, dysregulated NFAT3 signaling, and aberrant cell-cycle activity. MRE stiffening also drove fibroblast-to-myofibroblast transition in primary cardiac fibroblasts, recapitulating early fibrotic remodeling. Together, these findings position dynamically tunable MRE substrates as a human-relevant platform for dissecting mechano-genetic disease mechanisms and evaluating therapies targeting mechanotransduction pathways in HCM.

## Introduction

Hypertrophic cardiomyopathy (HCM) is the most common inherited cardiac disease, with an estimated prevalence ranging from 1 in 200 to 1 in 500 individuals (1, 2). Despite its genetic basis, HCM shows striking clinical heterogeneity: patients with HCM may remain asymptomatic, or develop heart failure, life-threatening arrhythmias, and sudden cardiac death (SCD), particularly among young adults (3–7). HCM is primarily caused by mutations in genes encoding sarcomeric proteins (8), most frequently in *MYBPC3* (myosin-binding protein C) and *MYH7* (β-myosin heavy chain) (9–11).

Cardiac hypertrophy exists on a spectrum: physiological hypertrophy is thought to be an adaptive, reversible response to stimuli such as postnatal growth, exercise, or pregnancy, whereas pathological hypertrophy is thought to be a maladaptive process driven by persistent mechanical stress such as chronic pressure overload (12–14). The latter is characterized by activation of hypertrophic signaling pathways that promote adverse structural remodeling, including left ventricular hypertrophy, diastolic dysfunction, interstitial fibrosis, and ultimately may progress to contractile dysfunction, arrhythmogenesis, and heart failure (13, 15–21). Historical evidence suggests that even for individuals harboring sarcomeric genomic variants linked to HCM, hypertrophic remodeling also falls on a spectrum: some genotype-positive individuals do not develop the phenotype (variable penetrance), and among those that do show the HCM phenotype, symptoms, location and severity of left ventricular hypertrophy are variable between individuals (variable expression) (22). Even within one family, in whom family members carry the same pathogenic variant, there is variable penetrance and expressivity (23, 24).

Incomplete penetrance and variable expression of HCM point to a potential role for non-genetic factors, such as mechanical stress, in shaping disease manifestation (25–30). Epidemiological evidence supports the hypothesis that blood pressure and the hemodynamic environment can modify HCM susceptibility and/or disease severity. Hypertension is a common condition, existing in approximately >40% of adults, regardless of whether or not they have HCM (31). Mendelian randomization studies provide support the influence of blood pressure in HCM: higher diastolic blood pressure was associated with sarcomere-negative HCM, defined as HCM without an identified pathogenic or likely pathogenic sarcomere-gene variant (32), whereas subsequent cross-ancestry analyses implicated higher systolic blood pressure in HCM overall (33). Large-scale genetic analyses have also implicated increased left ventricular contractility in both obstructive and nonobstructive HCM (34). Together, these findings support a potential contribution of blood pressure to HCM susceptibility. However, it remains unclear whether elevated blood pressure or the associated mechanical afterload modifies disease expression in carriers of pathogenic sarcomere variants.

Animal models have allowed the potential role of mechanical stress on HCM pathogenesis to be studied in a controlled and systematic manner. Studies that combined surgically induced mechanical afterload with a myocardium carrying a known sarcomere mutation support the notion that genetics and mechanics synergize together in HCM. In some cases, surgically induced afterload was added explicitly because mice harboring HCM-associated variants showed no overt disease pathology without this intervention. For example, several heterozygous *MYBPC3* mouse models exhibit little or no overt HCM phenotype under baseline conditions, supporting the use of secondary stressors to reveal disease phenotypes that are subtle or absent under baseline conditions (35, 36). In mice heterozygous for *MYH7* Arg403Gln mutation or for a *MYBPC3* truncating variant, transverse aortic constriction (TAC) produced greater hypertrophy and contractile dysfunction than wild-type mice subjected to the same pressure overload (35, 37).

Beyond revealing disease phenotypes in sarcomere-mutant animal models, TAC has been invaluable in demonstrating that mechanical afterload activates key hypertrophic signaling pathways, including MAPK (38) and calcineurin-NFAT (39), and accelerates disease progression even in the absence of sarcomeric mutations (40–42). However, this reliance on *in vivo* models, especially those that incorporate surgical maneuvers, carries significant limitations: these studies are costly, labor-intensive, user-skill-dependent, and prone to variable disease progression. More fundamentally, it is difficult to decouple the direct effects of mechanical loading from surgically induced inflammation (43). Finally, rodent models cannot recapitulate the specific genetic backgrounds and cardiomyocyte biology of human HCM (44).

Recent advances in human induced pluripotent stem cell (hiPSC) and genome editing technologies have opened a powerful alternative, enabling the study of HCM-causing mutations directly in cultured human cardiomyocytes (45). Patient-derived and isogenic hiPSC-cardiomyocytes (hiPS-CMs) have successfully recapitulated key disease hallmarks, including cellular hypertrophy, sarcomere disarray, hypercontractility, impaired relaxation, and dysregulated calcium cycling (46–52). Building on this, engineered 3D cardiac tissue models, developed by several groups, have allowed controlled application of mechanical forces simulating preload and afterload, further advancing the physiological relevance of these platforms (45, 53–58). Yet a fundamental limitation persists: most existing HCM models rely on substrates with fixed mechanical properties, presenting cells with a single static stiffness throughout the experiment.

This is a critical shortcoming, because the mechanical environment of the heart is inherently dynamic. *In vivo*, myocardial stiffness changes during development, across the cardiac cycle, and pathologically, as ECM remodeling following events such as myocardial infarction progressively increase tissue stiffness over time (59–61). In pressure-overload disease, this shift in mechanical loading drives the transition from adaptive to maladaptive remodeling (62–64). Static materials are insufficient to model this process, regardless of the stiffness value chosen (65, 66). What is needed is a platform that can dynamically and controllably stiffen in real time, within the same sample, to expose cardiomyocytes to a physiologically relevant progression of mechanical loading. The emergence of dynamically tunable biomaterials directly addresses this need. Platforms based on shape-memory polymers (60, 67–70), photo-programmable elastomers (71), and magnetorheological elastomers (MREs) (66, 68, 72) now enable real-time, reversible modulation of substrate stiffness. Among these, MREs have begun to demonstrate considerable promise for investigating cardiac mechanobiology: stiffness can be tuned rapidly and non-invasively by applying an external magnetic field, without chemical modification of the substrate or disruption of the cell culture. Prior work has demonstrated that dynamic mechanical cues delivered via MRE regulate cardiomyocyte behaviors including sarcomere reorganization and load-dependent functional adaptation, as well as fibroblast activation and mechanical memory effects that are simply inaccessible with static substrates (60, 68, 70).

However, whether MRE-based dynamic stiffening can unmask genotype-specific vulnerabilities in human HCM cardiomyocytes, and whether it can model the mechano-genetic interactions that drive early disease, is largely unexplored. Ramachandran *et al*. previously showed that iPS-CMs with a patient-specific *MYBPC3* premature STOP codon variant responded more aggressively to magnetically induced loading on MRE (73). These findings provide initial evidence that dynamic mechanical loading can interact with HCM-associated mutations to drive disease phenotype. However, molecular mechanisms linking the combination of mechanical loading and genetics to HCM phenotypes remain unclear.

In this study, we developed and characterized a tunable MRE platform to investigate how dynamically increased substrate stiffness interacts with a pathogenic *MYBPC3* frameshift mutation (*MYBPC3* <u>G1206</u>fs, *MYBPC3*fs (74, 75)) to drive early HCM-associated remodeling in hiPS-CMs. Leveraging this platform in concert with AI-driven automated single cell segmentation, we explored this pathological remodeling on the single-cell level and demonstrated that mechanical stiffening selectively triggers cellular pathology in *MYBPC3*fs cardiomyocytes, leading to sarcomere disorganization, increased expression of Nuclear Factor of Activated T-cells (NFAT3), one of the key transcriptional effectors of hypertrophic remodeling, and slightly elevated cell-cycle activity.

## Results

### Spatial dependence of magnetically-induced stiffening

Because hiPS-CMs beat at approximately 1 Hz, the substrate stiffness that cells work against is the dynamic modulus at that frequency. We therefore characterized the lateral extent of magnetically induced stiffening by spherical indentation of 50 wt% MREs at 1 mm increments of distance between the indentation site and the center of a neodymium magnet (**Fig. S1 A**), recovering the viscoelastic relaxation spectrum at each site and evaluating the dynamic reduced modulus |*M*^*^| (Methods). Although magnetic field strength fell off away from the magnet center (**Fig. S1B**), the relative reduction in stiffening was within measurement error (**Fig S1C**) until the distance from the magnet center exceeded 3mm. The loss tangent was likewise higher near the magnet (median 0.077 vs. 0.044; **Fig. S1D**), so magnetization increased both the stiffness and the dissipation experienced by cells at beating frequency, although tan*δ* remained below 0.1 throughout and the substrate was therefore predominantly elastic at 1 Hz. Because the contact diameter (approximately 2.7 mm) exceeded the spacing between indentation sites, neighboring measurements were not independent, and we therefore compare pooled regions rather than a site-by-site profile. Based on these findings, we generated 6.4 mm-diameter stencils to confine seeded hiPS-CMs or human primary ventricular cardiac fibroblasts (cFBs) within 3.2 mm of the center of the magnet, ensuring that all cells experienced similar magnet-induced substrate stiffening in subsequent experiments (**Fig. S1 F, G**).

### Increasing CIP wt% increases MRE stiffness while maintaining cytocompatibility

To broaden the range of stiffness achieved when applying the magnetic field, we tested whether changing the CIP content of MREs changed their mechanical properties (76, 77). As accurately sampling the compressive mechanical properties with materials like PDMS, which stick to test probes and thus often prevent appropriate lateral expansion (Poisson effect) is challenging (78), and because adherent cell monolayers work mechanically against their substrates primarily in shear (79), we applied a static shear mechanical test devised previously (68) to measure MRE stiffness either with or without the magnet applied.

Under unmagnetized conditions (baseline), shear modulus did *not* differ significantly between 33-66 wt% MREs from unfilled 0 wt% elastomer, whereas increasing CIP wt% to 75 or higher led to a significant baseline shift (**Fig. 1A**). However, upon magnetization, 66 wt% substrates showed significantly higher shear modulus than 50 wt% substrates (**Fig. 1B-D**). Consistent with this, the stiffness magnetorheological (SMR) effect, which describes the percent increase in magnet-induced MRE stiffness, was higher at 66 wt% (**Fig. 1E**). Together, these findings indicate that increasing CIP content to 66 wt% maximizes the range of stiffness over which MRE can be actuated magnetically, without altering the baseline rigidity. To ensure that higher CIP content did not introduce cytotoxicity, we assessed cell viability using an Alamar Blue assay. C2C12 cells exposed to MRE-conditioned media showed no reduction in metabolic activity at CIP contents up to 66 wt%, whereas media conditioned with non-encapsulated CIP particles (100 wt%) caused a significant loss of viability (**Fig. 1F**). Together with the mechanical testing results, these findings identify 66 wt% CIP loading (2.95 mm thickness) as the formulation that achieves the highest shear modulus while maintaining cytocompatibility, making it suitable for downstream biological studies.

**Figure 1.**
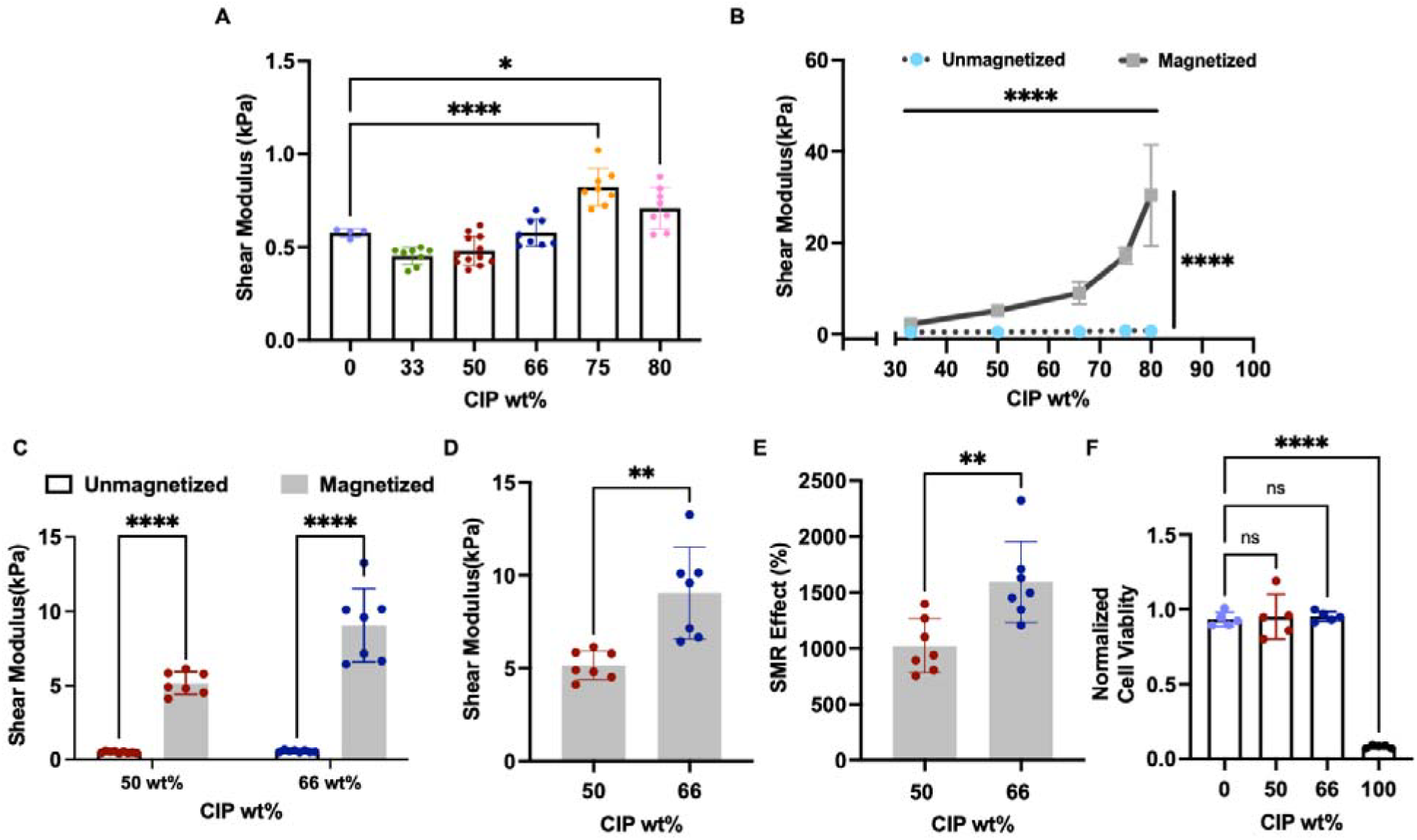
Increasing CIP content enhances the magnetically tunable stiffness of MREs while maintaining cytocompatibility. **(A)** Baseline shear modulus of unmagnetized MREs containing increasing CIP content (0-80 wt%), showing comparable baseline stiffness up to 66 wt% CIP, with significantly higher stiffness observed at ≥75 wt%. **(B)** Shear modulus of unmagnetized and magnetized MREs across increasing CIP content (33-80 wt%) demonstrate progressively greater magnetically induced stiffening with increasing CIP loading. Quantification of **(C)** shear modulus of 50 wt% and 66 wt% MREs under unmagnetized and magnetized conditions, **(D)** magnetized shear modulus, and **(E)** SMR effect (%), comparing 50 wt% and 66 wt% formulations. Increasing CIP loading from 50 wt% to 66 wt% significantly enhanced magnetically-induced stiffness and SMR effect while maintaining comparable baseline stiffness. **(F)** Normalized cell viability of C2C12 cells cultured in media conditioned with MREs containing increasing CIP content. Cell viability was maintained for formulations containing up to 66 wt% CIP, whereas exposure to non-encapsulated CIP particles (100 wt%) significantly reduced viability. Data are presented as mean ±*SD*; *P* values were determined using (**A, F**) one-way ANOVA with Dunnett correction for multiple comparison, (**B, C**) two-way ANOVA with Holm-Šidák for multiple comparisons, and (**D, E**) unpaired t-test Welch’s correction; **** *p* < 0.0001, *** *p* < 0.001, ** *p* < 0.01, * *p* < 0.05, ns = not significant.

### Mechanical stimulation induces fibroblast-to-myofibroblast transition

Because cardiac fibroblasts are key mechanosensitive regulators of fibrotic remodeling, we next assessed the impact of MRE stiffening on cFB activation (**Fig. 2A**). Automated cell segmentation using the open-source CellPose software package (Cyto2 model; Supplementary Section 1.2) revealed that cFB on magnetized substrates showed a significantly decreased cell area compared to both unmagnetized controls and TGF-β-treated cells (**Fig. 2B, C**). In contrast, no significant differences were observed in cell eccentricity (**Fig. 2D**), elongation (**Fig. S3 A**), circularity (**Fig. S3 B**), or cell density per region of interest (**Fig. S3 C**) across conditions, indicating that mechanical stimulation selectively affects cell size without broadly altering cell shape.

**Figure 2.**
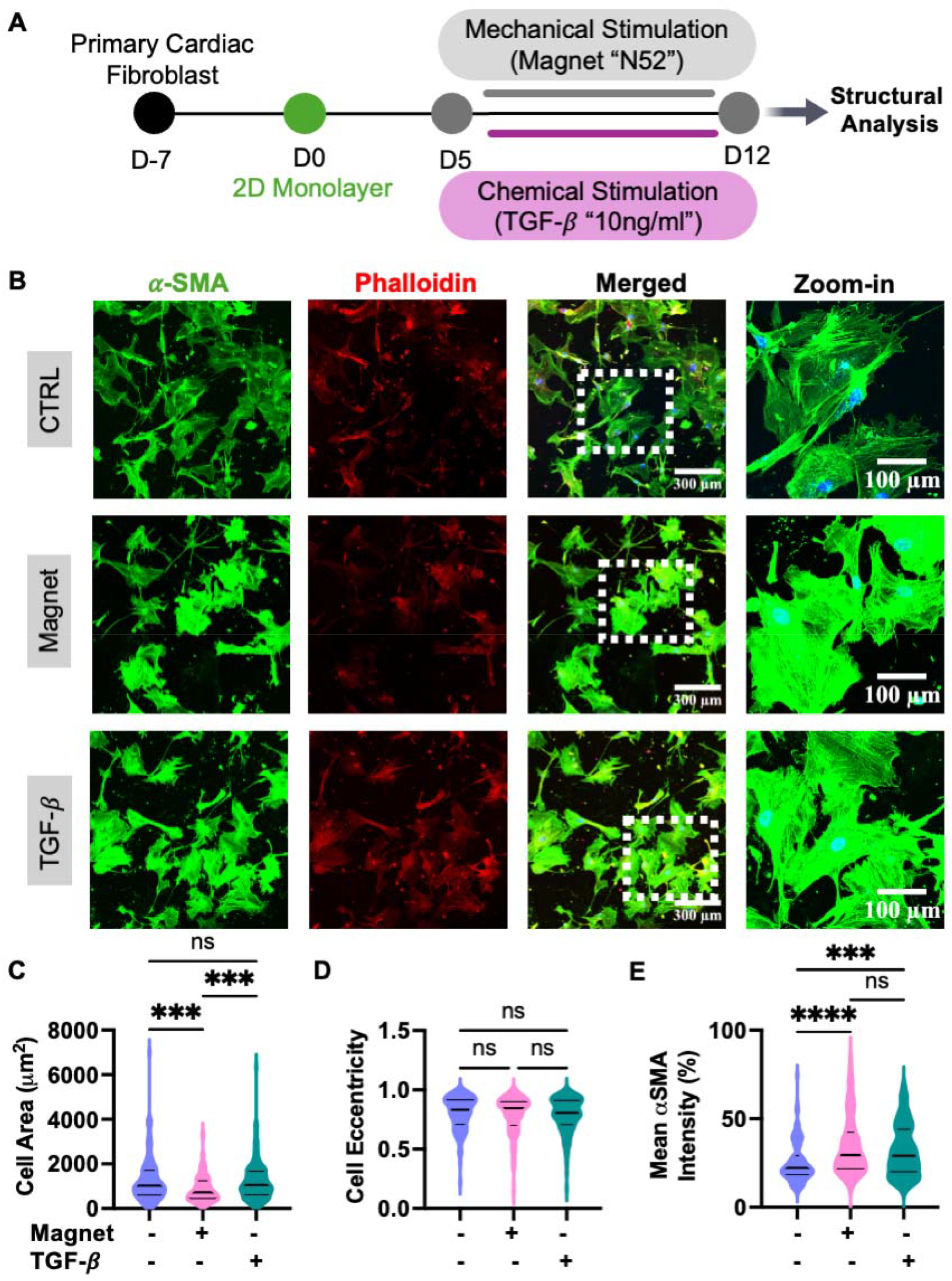
Mechanical and chemical stimulation regulate cFB morphology and activation. **(A)** Schematic of the experimental timeline. cFB were seeded as a 2D monolayer with 25k/cm^2^ density on day 0, followed by application of mechanical stimulation (Magnet “N52”) or chemical stimulation (TGF-*β*, 10 ng/ml) from day 5 to day 12, after which structural analysis was performed. **(B)** Representative immunofluorescence images of cardiac fibroblasts under three conditions; control (CTRL), mechanical stimulation (Magnet), and chemical stimulation (TGF-*β*) — stained for *α*-SMA (green), Phalloidin (red), and Hoechst (blue), shown as individual channels, merged, and zoomed-in views. Scale bars: 300 *µ*m (merged) and 100 *µ*m (zoom-in). Quantification of **(C)** cell area, **(D)** cell eccentricity, and **(E)** mean *α*-SMA intensity per cell (%) across CTRL, Magnet, and TGF-*p* conditions. Mechanical stimulation did increase *α*-SMA levels comparable to TGF-*p* treated condition. Cell eccentricity was not significantly different across any condition. Data are presented as violin plots, n represent individual cells; *P* values by one-way ANOVA, non-parametric, Kruskal-Wallis test with Dunn’s multiple comparisons; **** *p* < 0.0001, *** *p* < 0.001, ns = not significant.

Importantly, despite the reduction in cell area, expression of α-SMA, a marker of fibroblast-to-myofibroblast transition (80), was significantly elevated in magnetized cFB relative to unmagnetized controls and reached levels comparable to those induced by TGF-β treatment (**Fig. 2E**). This myofibroblast activation suggests that MRE-based mechanical stimulation engages fibrotic signaling pathways independently of gross cell spreading. These findings were consistent across seeding densities (10,000 and 25,000 cells/cm^2^; **Fig. 2; Fig. S4**), suggesting that changes in cell-cell contact were not a dominant factor in cFB activation. Together, these data demonstrate that mechanical stimulation alone is sufficient to drive fibroblast-to-myofibroblast transition, paralleling the fibrotic activation induced biochemically by TGF-β and recapitulating mechanical activation pathways relevant to early HCM progression and post-MI remodeling (72, 81).

### *MYBPC3*fs cardiomyocytes are more susceptible to pathological structural remodeling under mechanical loading

Having established a mechanically tunable MRE platform, we next investigated the response of isogenic control (WT) and *MYBPC3*fs hiPS-CM monolayers to the same mechanical stimulus. Following 1 week of magnetization (**Fig. 3A**), isogenic control hiPS-CMs showed no significant change in cell area upon magnetization (**Fig. 3B; Fig. S5 A**). Structural analysis revealed a modest but significant decrease in cell circularity upon magnetization (**Fig. S5 B**), whereas cell elongation and eccentricity remained unchanged (**Fig S5 C, D**). These findings of modest morphological changes in healthy cardiomyocytes suggest that they largely tolerate increased mechanical loading without undergoing overt pathological remodeling.

**Figure 3.**
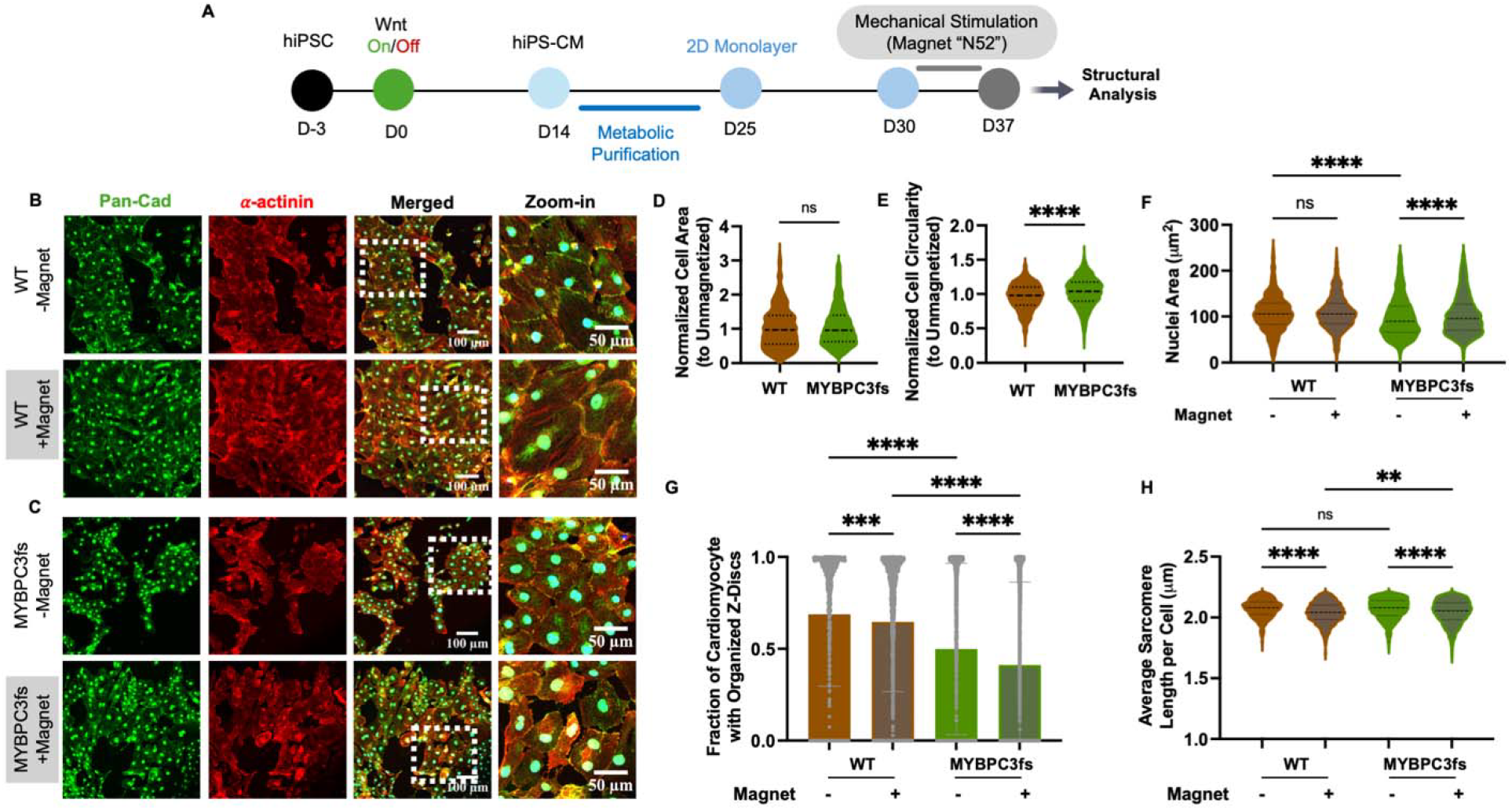
*MYBPC3*fs hiPSC-cardiomyocytes show hypertrophic remodeling and sarcomere disorganization in response to mechanical stimulation. **(A)** Schematic of the experimental timeline. hiPSCs underwent Wnt-modulated differentiation into hiPS-CMs beginning at day 0, followed by metabolic purification (day 14-25), monolayer formation (day 25), and 7 days of mechanical stimulation via Magnet “N52” (day 30–37), after which structural analysis was performed. **(B, C)** Representative immunofluorescence images of isogenic control (WT) **(B)** and *MYBPC3*fs **(C)** hiPS-CMs under unmagnetized (−Magnet) and magnetized (+Magnet) conditions, stained for Pan-Cadherin (green), *α*-actinin (red), and Hoechst (blue), shown as individual channels, merged, and zoomed-in views. Scale bars: 100 *µ*m (merged) and 50 *µ*m (zoom-in). Quantification of **(D)** normalized cell area and **(E)** normalized cell circularity relative to their respective unmagnetized controls for WT and *MYBPC3*fs hiPS-CMs. Normalized cell circularity (to respective unmagnetized baseline) was significantly higher in *MYBPC3fs* than in WT cardiomyocytes, whereas normalized cell area did not differ significantly between genotypes. **(F)** Nuclear area across all four conditions. *MYBPC3*fs cardiomyocytes showed smaller baseline nuclear area compared to WT controls; magnetization increased nuclear area in *MYBPC3*fs cells. **(G)** Fraction of cardiomyocytes with organized Z-discs across all conditions. Magnetization reduced sarcomere organization in WT cardiomyocytes, and *MYBPC3fs* cells showed lower baseline sarcomere organization that was further reduced upon magnetization. **(H)** Average sarcomere length per cell, calculated from all Z-disc vectors in each cell. Sarcomere length was significantly reduced in both WT and *MYBPC3*fs cardiomyocytes upon magnetization, with *MYBPC3*fs cells showing lower length under magnetized conditions compared to magnetized WT cells. Data are presented as violin plots, n represent individual cells (**D, E, F, H**) or bar plots, n represent each component (**G**). *P* values in (**D, E**) by unpaired t-test, non-parametric, Mann-Whitney U test; in (**F, G, H**) by one-way ANOVA, non-parametric, Kruskal-Wallis test with Dunn’s multiple comparisons; **** *p* < 0.0001, *** *p* < 0.001, ** *p* < 0.01, ns = not significant.

In contrast to isogenic control hiPS-CMs, under the same mechanical conditions, cardiomyocytes harboring the *MYBPC3*fs mutation (75) exhibited more extreme structural remodeling in response to MRE magnetization, with a significant increase in both cell area and cell circularity (**Fig. 3C; Fig. S6 A, B**), while cell elongation and eccentricity remained unchanged (**Fig. S6 C, D**). This shift toward a larger, rounder cellular morphology is consistent with concentric hypertrophic growth in HCM (3).

To directly compare genotype-dependent responses to mechanically induced stiffening, we normalized the morphological response of magnetized cells to their respective unmagnetized controls for each genotype to quantify the effect of mechanical loading independent of baseline differences. Notably, while absolute cell area was significantly larger in magnetized *MYBPC3*fs cardiomyocytes compared to their unmagnetized baseline (**Fig. S6 A**), the normalized fold-change in cell area did not differ significantly between isogenic control and *MYBPC3*fs cardiomyocytes (**Fig. 3D**), likely because both genotypes showed a similar relative increase in cell area (median ~11%), which reached significance only in *MYBPC3f*s cardiomyocytes, whose cell-area distribution was narrower (**Fig. S5A; Fig. S6A**). Beyond cell area, magnetization produced a significantly greater increase in cell circularity in *MYBPC3*fs cardiomyocytes compared to isogenic controls (**Fig. 3E**), and *MYBPC3*fs cells showed a significantly greater reduction in normalized cell elongation upon magnetization compared to isogenic controls (**Fig. S7 A**). Normalized cell eccentricity did not differ significantly between genotypes (**Fig. S7 B**), indicating a more pronounced shift toward an isotropic, hypertrophic morphology in hiPS-CMs harboring the *MYBPC3* variant. Together, these data reveal that while both genotypes experience the same mechanical stimulus, *MYBPC3*fs cardiomyocytes undergo greater shape remodeling.

### *MYBPC3*fs cardiomyocytes have smaller nuclei at baseline and undergo mechanically-induced nuclear remodeling

In cardiomyocytes, mechanical forces are transmitted directly to the nucleus through the desmin-lamin cytoskeletal network, where they regulate nuclear architecture and mechanosensitive gene expression (82). Moreover, HCM-associated sarcomere variants have been linked to changes in nuclear morphology (83). Consistent with this prior observation, while magnetization did not alter nuclear area in isogenic control cells, nuclear area increased significantly in *MYBPC3*fs cardiomyocytes upon magnetization, while *MYBPC3*fs cardiomyocytes have consistently smaller nuclear area compared to isogenic control at baseline (**Fig. 3F**). These changes in nuclear size were accompanied by nuclear shape remodeling. In isogenic control cardiomyocytes, magnetization significantly increased nuclear circularity while nuclear elongation and eccentricity remained unchanged (**Fig. S8 A-C**). In contrast, within *MYBPC3*fs cardiomyocytes, magnetization significantly decreased nuclear eccentricity and elongation while increasing nuclear circularity, indicating a shift toward a rounder, more isotropic nuclear morphology upon mechanical stimulation (**Fig. S8 A-C**). Together, these findings suggest that *MYBPC3*fs cardiomyocytes not only start with structurally distinct nuclei at baseline (unmagnetized) but are also more responsive to mechanical stimulation at the nuclear level, pointing to a broader mechanosensitive remodeling response that extends beyond the cytoskeleton to the nucleus itself.

### *MYBPC3*fs cardiomyocytes show lower sarcomere organization at baseline that is further exacerbated by mechanical stimulation

Given that HCM frequently involves sarcomere disorganization, we next analyzed sarcomere structure under mechanical loading in both genotypes using a modified version of the SarcAsM algorithm developed by Härtter *et al*. (84). Sarcomere regions were classified as organized based on the following criteria: 1) length of 0.8-2.2 μm, 2) orientation standard deviation ≤ 20°, 3) ≥ 3 sarcomeres per region, 4) ≥ 3 sarcomere repeats per region, and 5) exclusion of regions with ≥ 10% saturated pixels (**Fig. S9**).

Quantification showed that magnetization significantly reduced the fraction of cardiomyocytes with organized Z-discs in isogenic controls (**Fig. 3G**), indicating that increased mechanical afterload alone is sufficient to partially disrupt sarcomere organization even in healthy cardiomyocytes. Consistent with the known structural instability associated with *MYBPC3* loss-of-function variants (35), unmagnetized *MYBPC3*fs cardiomyocytes showed a significantly lower fraction of organized sarcomere area compared to unmagnetized isogenic controls, and mechanical loading further exacerbated this deficit (**Fig. 3G**).

In prior HCM studies, a reduced resting sarcomere length has been observed, and was associated with impaired relaxation (85, 86). Thus, we used the SarcAsM package to assess resting sarcomere length. In isogenic control cardiomyocytes, magnetization significantly reduced sarcomere length compared to their unmagnetized baseline (**Fig. 3H**), a finding that is inconsistent with maturation (87) and instead suggests the possibility of mechanically driven impairment of diastolic relaxation (85). At baseline, sarcomere length was not significantly different between isogenic control and *MYBPC3*fs cardiomyocytes. However, upon magnetization, *MYBPC3*fs cardiomyocytes showed significantly lower sarcomere length than magnetized isogenic control cells (**Fig. 3H**). This suggests that the frameshift mutation makes hiPS-CMs more vulnerable to mechanical disruption of sarcomere architecture – likely leading to diastolic dysfunction. Together, these data reveal a genotype-dependent sensitivity to mechanical loading in which *MYBPC3*fs cardiomyocytes exhibit exaggerated sarcomere disorganization under identical mechanical conditions, recapitulating the structural vulnerability to pressure overload that is characteristic of *MYBPC3*-associated HCM in animal models settings (37, 73).

### Mechanical stimulation alters NFAT3 levels and subcellular distribution in hiPS-CMs harboring the HCM *MYBPC3*fs mutation

The calcineurin-NFAT signaling pathway plays a central role in pathological cardiac hypertrophy and is activated by increased mechanical load and altered calcium handling, both hallmark features of HCM (88). In response to elevated pressure overload, mechanosensitive channels open, leading to increased intracellular Ca^2+^, which activates calcineurin, enabling dephosphorylation and nuclear translocation of NFAT transcription factors and ultimately driving hypertrophic gene expression (**Fig. 4A**) (88). Aberrant or sustained NFAT activation is considered a key molecular driver of maladaptive remodeling in HCM (14).

**Figure 4.**
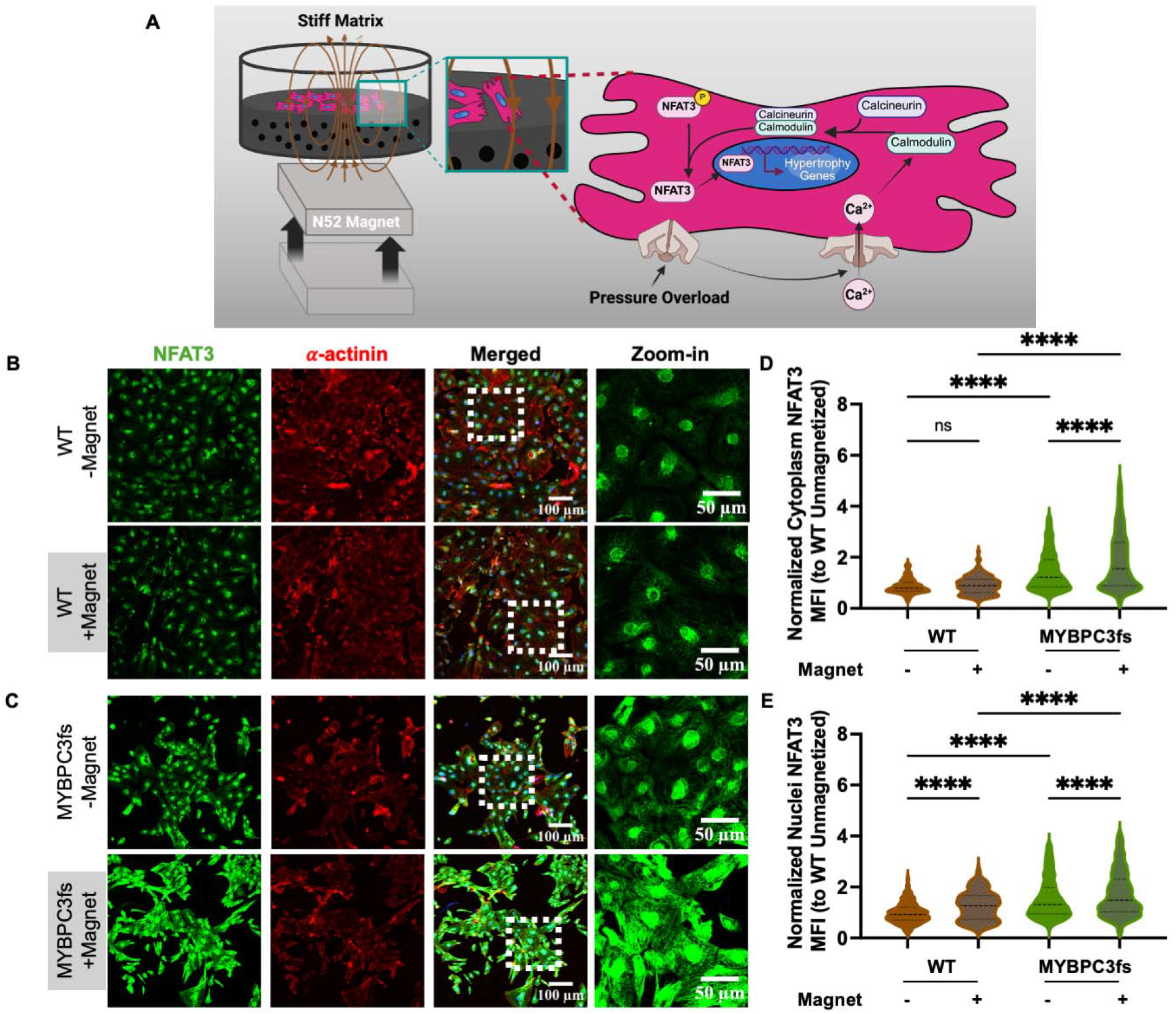
*MYBPC3*fs hiPS-CMs exhibit elevated NFAT3 levels that are further increased by mechanical stimulation. **(A)** Schematic illustrating the proposed mechanosensitive calcineurin-NFAT signaling pathway. Magnetization of MRE substrates increases substrate stiffness, simulating pressure overload, which elevates intracellular Ca^2+^, activates calcineurin-calmodulin, and drives dephosphorylation and nuclear translocation of NFAT3, ultimately upregulating hypertrophic gene expression. **(B, C)** Representative immunofluorescence images of **(B)** isogenic control and **(C)** *MYBPC3*fs hiPS-CMs under unmagnetized (−Magnet) and magnetized (+Magnet) conditions, stained for NFAT3 (green) and *α*-actinin (red), shown as individual channels, merged, and zoomed-in views. Scale bars: 100 *µ*m (merged) and 50 *µ*m (zoom-in). Quantification of **(D)** normalized cytoplasmic NFAT3 mean fluorescence intensity (MFI) and **(E)** normalized nuclear NFAT3 MFI, both normalized to the isogenic control unmagnetized condition. Magnetization did not significantly alter cytoplasmic NFAT3 levels but increased nuclear levels in isogenic control cardiomyocytes, whereas *MYBPC3*fs cardiomyocytes showed a robust and significant increase in both cytoplasmic and nuclear NFAT3 MFI upon magnetization. Data are presented as violin plots, n represent individual cell; *P* values by one-way ANOVA, non-parametric, Kruskal-Wallis test with Dunn’s multiple comparisons; **** *p* < 0.0001, ns = not significant.

To investigate whether MRE-based substrate stiffening activates this mechanosensitive pathway, we assessed the overall expression and subcellular localization of NFAT3 in isogenic control and *MYBPC3*fs hiPS-CMs before and after magnetization. At baseline, isogenic control and *MYBPC3*fs cardiomyocytes showed comparable nuclear-to-cytoplasmic (nuc/cyto) NFAT3 ratios (**Fig. S10 A**). Following magnetization, isogenic control cardiomyocytes showed a significant increase in the nuc/cyto ratio (**Fig. S10 A**), suggesting active NFAT3 nuclear translocation in response to mechanical stimulation. In contrast, *MYBPC3*fs cardiomyocytes showed a significant decrease in the nuc/cyto ratio upon magnetization (**Fig. S10 A**), suggesting a redistribution of NFAT3 toward the cytoplasm in the mutant line. Importantly, when normalized to their respective unmagnetized baselines, *MYBPC3*fs cardiomyocytes showed a significantly lower relative NFAT3 response compared to isogenic control upon magnetization (**Fig. S10 B**).

While relative NFAT3 translocation may have been limited in the HCM cardiomyocytes, overall NFAT3-driven transcription ultimately depends on the total amount of this protein in the nucleus. Thus, we assessed overall cellular NFAT3 expression. Quantification of absolute cytoplasmic and nuclear NFAT3 mean fluorescence intensity (MFI) (normalized to unmagnetized isogenic control) at baseline showed that unmagnetized *MYBPC3*fs cardiomyocytes had significantly higher cytoplasmic and nuclear NFAT3 levels compared to unmagnetized isogenic controls (**Fig. 4B-E**). Upon magnetization, both cytoplasmic and nuclear NFAT3 MFI increased significantly in *MYBPC3*fs cardiomyocytes (**Fig. 4B-E**). In contrast, isogenic control cardiomyocytes showed no significant change in cytoplasmic NFAT3 MFI after chronic magnetization, while nuclear NFAT3 MFI increased significantly. The observed decrease in the nuc/cyto ratio in magnetized *MYBPC3*fs cardiomyocytes (**Fig. S10 A**), despite the significant increase in absolute nuclear NFAT3 levels, may suggest that the nucleus may have reached a saturation point in its capacity to accommodate further NFAT3 translocation in the *MYBPC3*fs cardiomyocytes, so that additional NFAT3 accumulates in the cytoplasm. In this context, the disproportionate accumulation of NFAT3 in the cytoplasm relative to the nucleus upon magnetization would reduce the nuc/cyto ratio even as total and nuclear NFAT3 levels rise. In turn, this suggests that the *MYBPC3*fs line operates under chronically elevated NFAT3 signaling that becomes overwhelmed rather than further amplified upon additional mechanical challenge.

### *MYBPC3*fs cardiomyocytes exhibit an altered cell-cycle response to mechanical stimulation

A key observation from our morphological analysis was that unmagnetized *MYBPC3*fs cardiomyocytes had significantly smaller cell area compared to isogenic controls (**Fig. S5 A** and **Fig. S6 A**), which is seemingly at odds with the hypertrophic phenotype expected in HCM. We hypothesized that rather than hypertrophic growth of individual cells, the *MYBPC3*fs mutation may promote hyperplastic growth, in which increased cardiomyocyte number rather than increased cell size contributes to cardiac enlargement. Such a phenomenon has been observed in postnatal rodent hearts (89, 90). To test for potential cell cycle changes in our hiPS-CMs, we assessed Ki67 expression (**Fig. 5A, B**). Ki67 positivity was calculated as the number of Ki67-positive nuclei divided by the total number of Hoechst-stained nuclei per ROI.

**Figure 5.**
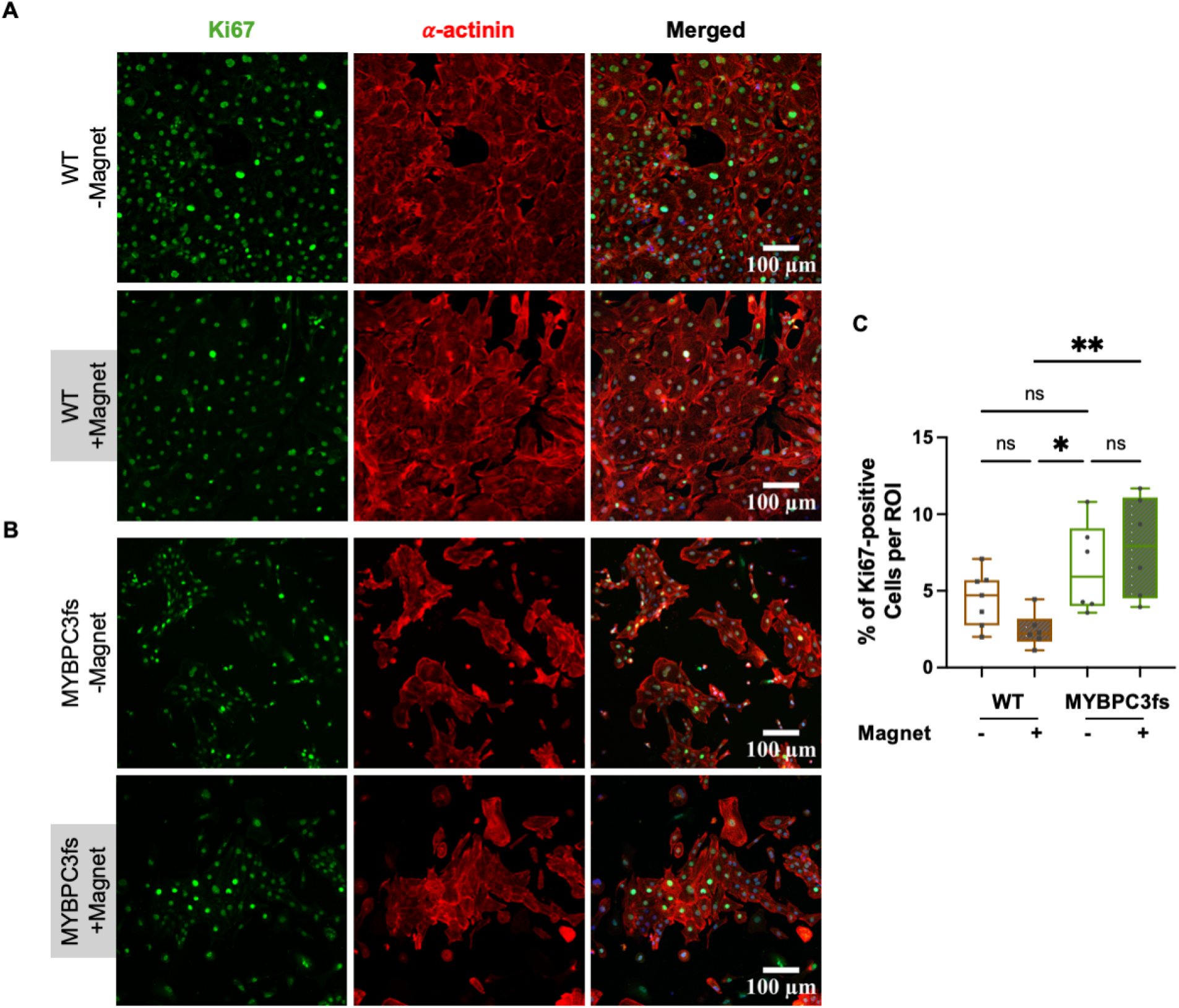
*MYBPC3*fs hiPS-CMs show greater Ki67 positivity than isogenic controls under mechanical stimulation. **(A, B)** Representative immunofluorescence images of **(A)** isogenic control and **(B)** *MYBPC3*fs hiPS-CMs under unmagnetized (−Magnet) and magnetized (+Magnet) conditions, stained for Ki67 (green), *α*-actinin (red), and Hoechst (blue), shown as individual channels and merged. Scale bars: 100 *µ*m. **(C)** Quantification of the percentage of Ki67-positive cells per region of interest (ROI) across all four conditions. At baseline, *MYBPC3*fs cardiomyocytes showed a trend toward higher proportion of Ki67-positive cells compared to unmagnetized isogenic controls, indicating a slightly elevated cell cycle activity in the mutant line. Magnetization resulted in a trend toward lower Ki67 positivity in isogenic control cardiomyocytes, whereas a trend toward increased proliferation was observed in magnetized *MYBPC3*fs cells, though this did not reach statistical significance. However, Ki67 positivity was significantly higher in magnetized *MYBPC3*fs cells than in magnetized isogenic control cardiomyocytes. Data are presented as box plots, n represent individual ROI; *P* values by one-way ANOVA, parametric, Holm-Šidák for multiple comparisons; * *p* < 0.05, ns = not significant.

Quantification showed that unmagnetized *MYBPC3*fs cardiomyocytes had a trend toward higher percentage of Ki67-positive cells per ROI compared to unmagnetized isogenic controls, though not significant (**Fig. 5C**), indicating a slightly elevated baseline cell cycle activity in the mutant line. Interestingly, magnetization led to a non-significant trend toward fewer Ki67-positive cells in isogenic control cardiomyocytes (**Fig. 5C**), suggesting that increased mechanical afterload may suppresses cell cycle activity in healthy cardiomyocytes, consistent with the shift toward physiologic hypertrophic rather than hyperplastic growth under pressure overload. In contrast, magnetization did not significantly alter Ki67 positivity in *MYBPC3*fs cardiomyocytes, while it was significantly elevated in magnetized *MYBPC3*fs cells compared to magnetized isogenic controls (**Fig. 5C**). These findings suggest that the smaller baseline cell size observed in *MYBPC3*fs cardiomyocytes may reflect an actively proliferating population that has not yet undergone hypertrophic growth, and that slightly elevated cell cycle activity in hiPS-CM harboring the HCM variant may represent an early disease mechanism that precedes and contributes to pathological cardiac remodeling in *MYBPC3*-associated HCM. Together, these data are consistent with a model in which dysregulated cell cycle activity and elevated proliferation precede and potentially start the hypertrophy in early *MYBPC3*-associated HCM, where increased cardiomyocyte number rather than increased cell size may be the primary contributor to cardiac enlargement at early disease stages (91, 92).

## Discussion

In this study, we developed and applied a dynamically tunable magnetorheological elastomer (MRE) platform to model the mechano-genetic interactions that drive early hypertrophic cardiomyopathy (HCM). Under the same mechanical stimulus that simulates pressure overload, isogenic control and *MYBPC3*fs hiPS-CMs showed distinct structural and signaling responses. We demonstrated that the patient-specific *MYBPC3*fs genomic variant increases the susceptibility of human cardiomyocytes to mechanical stress, resulting in exaggerated hypertrophic remodeling, sarcomere disorganization, altered nuclear remodeling, dysregulated NFAT3 signaling, and slightly elevated cell-cycle activity. Together, these findings demonstrate that the response of human cardiomyocytes to mechanical loading is influenced by sarcomeric genomic variants. Our findings further establish MRE-based dynamic stiffening as a human-relevant platform for investigating genotype-dependent mechano-genetic interactions in cardiomyopathy.

Although pathogenic sarcomeric variants are established causes of HCM, the incomplete penetrance and variable severity of this disease suggest that the primary mutation alone may not fully predict disease onset or severity (22, 30). This is particularly evident in *MYBPC3*-associated HCM, in which carriers of identical truncating variants exhibit broad variability in maximal wall thickness, and common genetic variation further modifies phenotypic severity (32, 93). Variable disease expression in HCM is perhaps best demonstrated by monozygotic twins carrying the same pathogenic sarcomere variants, who can develop discordant left ventricular hypertrophy despite nearly identical genetic background (23).

Recent human genetic studies incorporating Mendelian randomization analyses have implicated elevated blood pressure in HCM susceptibility, supporting a potential role for hemodynamic loading as a modifiable contributor to disease expression (32–34). However, the blood-pressure association stratified by sarcomere status was demonstrated primarily in sarcomere-negative HCM (32), whereas subsequent studies evaluated all-comer or obstructive and nonobstructive HCM without resolving the effect of blood pressure in pathogenic sarcomere-variant carriers (33, 34). Thus, whether increased mechanical loading that simulates pressure overload modifies disease severity in sarcomere-positive HCM remains incompletely understood (94).

Controlled animal and engineered human cardiac models have begun to address the underlying mechanisms through which increased mechanical resistance to heart contraction might exacerbate pathological remodeling in the myocardium of individuals harboring sarcomeric gene-mutations (65, 95–99). Notably, recent dynamic biomaterial studies demonstrated that mechanical loading can reveal *MYBPC3*-dependent sarcomere abnormalities that are less apparent under static conditions (65). Consistent with these observations, our findings show that *MYBPC3*fs cardiomyocytes exhibit exaggerated structural and signaling responses to the same mechanical stimulus compared with isogenic controls, providing evidence that sarcomeric variant influences cardiomyocyte adaptation to mechanical loading (65).

At the structural level, dynamic MRE-substrate stiffening induced distinct remodeling responses in *MYBPC3*fs and isogenic control hiPS-CMs. Mechanical afterload increased cell area and circularity in *MYBPC3*fs cardiomyocytes, whereas control cells showed no increase in cell area and a reduction in circularity, indicating that sarcomeric genetic variant alters the direction of cellular adaptation to the same mechanical stimulus. A similar genotype-dependent response was observed at the nuclear level, where stiffening increased nuclear area selectively in *MYBPC3*fs cardiomyocytes and produced a more pronounced shift toward rounder nuclei (lower elongation and eccentricity) in *MYBPC3*fs than in isogenic control cardiomyocytes. *MYBPC3*fs cardiomyocytes also exhibited reduced sarcomere organization under baseline conditions, which was further impaired following mechanical stiffening. In human HCM myocardium, myofiber disarray is more prominent in patients carrying pathogenic or likely pathogenic sarcomere variants and has been proposed as an early feature of disease (100). Mechanical afterload also reduced resting sarcomere length in both genotypes, with the shortest sarcomere length observed in magnetized *MYBPC3*fs cardiomyocytes. Because cardiomyocyte maturation is generally accompanied by *increased* sarcomere length (87), this reduction is unlikely to reflect enhanced maturation. In micropatterned patient-specific HCM iPS-CMs, Wu *et al*. similarly reported shorter diastolic sarcomere length across *MYH7*-, *MYBPC3*-, and *TNNT2*-mutant cells, accompanied by slower relaxation velocity and prolonged relaxation duration (86). Therefore, we hypothesize that this aberrant mechanosensitivity is likely rooted in the structural consequences of *MYBPC3* loss-of-function variants: reduced cMyBP-C levels compromise the mechanical stability of the sarcomere and impair its ability to buffer mechanical stress, lowering the threshold at which mechanical loading triggers maladaptive remodeling (35).

A key and unexpected finding of this study was that unmagnetized *MYBPC3*fs cardiomyocytes were actually smaller than isogenic controls at baseline, rather than hypertrophied as might be expected. This finding mirrors previous reports by others studying HCM in zebrafish (101), and likely the observation is enabled by the relative developmental immaturity of iPS-CM. Consistent with our observation, *MYBPC3*fs cardiomyocytes showed non-significant trend toward elevated Ki67 positivity at baseline and significantly higher Ki67 positivity than isogenic control upon magnetization, indicating increased cell cycle activity. This is particularly significant because cell-cycle activity is limited in differentiated hiPS-CMs, despite their relative developmental immaturity (87). Interestingly, re-entry into the cell cycle is a recognized feature of pathological cellular remodeling in postnatal cardiomyocytes (85, 90–92). Human genetics have also identified an HCM-associated locus near *CDKN1A*, which encodes the cyclin-dependent kinase inhibitor p21, implicating cell-cycle regulation in HCM susceptibility (32).

Importantly, mechanical stimulation led to a trend toward lower cell cycle activity in isogenic control cardiomyocytes, consistent with the known shift from hyperplastic to hypertrophic growth under pressure overload (102, 103), while *MYBPC3*fs cardiomyocytes failed to suppress proliferation under the same mechanical conditions, where Ki67 positivity was significantly higher in *MYBPC3*fs cells than in isogenic control cardiomyocytes. Therefore, we propose that MYBPC3fs cardiomyocytes may favor hyperplasia rather than hypertrophic growth at early disease stages. This dysregulated mechano-proliferative response raises the possibility that in early *MYBPC3*-associated HCM, increased cardiomyocyte number rather than increased cell size may be the primary contributor to cardiac enlargement, and that hypertrophic growth may emerge only later as the disease progresses (89, 90). This concept has important implications for disease staging and the interpretation of cardiac imaging findings in early HCM, where wall thickening may precede detectable cellular hypertrophy.

Consistent with the known role of calcineurin-NFAT signaling in pathological hypertrophy from clinical HCM tissue biopsies (104), we found that MRE stiffening activated NFAT3 in a genotype-dependent manner. Unexpectedly however, while isogenic control cardiomyocytes showed a clear increase in the nuclear-to-cytoplasmic NFAT3 ratio upon magnetization, indicative of active nuclear translocation, *MYBPC3*fs cardiomyocytes showed a decrease in this ratio despite a significant increase in absolute nuclear and cytoplasmic NFAT3 levels. We interpret this as potential evidence that *MYBPC3*fs cardiomyocytes may operate under chronically elevated baseline NFAT3 activity, such that the nucleus becomes saturated in its capacity to accommodate further NFAT3 upon additional mechanical challenge, resulting in cytoplasmic accumulation and a reduced nuc/cyto ratio. This model is consistent with prior reports that *MYBPC3*-deficient cardiomyocytes exhibit elevated myofilament Ca^2+^ sensitivity and dysregulated calcium handling (105), which would keep the calcineurin-NFAT pathway chronically active even without additional mechanical stress (99). Finsen *et al*. further demonstrated that cyclic stretch activates calcineurin-NFAT signaling in cardiomyocytes through the Z-disc/costamere mechanotransducer syndecan-4 (106). The combination of chronic NFAT activation and mechanical stiffening may therefore represent a two-way mechanism that accelerates pathological gene reprogramming in mutation-positive HCM (88, 107).

Beyond cardiomyocytes, our data suggest that MRE-based mechanical stimulation alone is sufficient to drive fibroblast-to-myofibroblast transition in primary cardiac fibroblasts, evidenced by significantly elevated α-SMA expression in magnetized cells that was comparable to the biochemical positive control TGF-β. This finding is particularly relevant to HCM pathophysiology, where interstitial fibrosis driven by myofibroblast activation is a key determinant of diastolic dysfunction and disease severity (72, 81). The ability of our platform to model both cardiomyocyte hypertrophic remodeling and fibroblast activation in parallel positions it as a versatile integrated tool for studying the multicellular mechanobiology of HCM.

Compared to existing *in vitro* systems, our MRE-based approach offers several conceptual and practical advantages. Most critically, the ability to dynamically modulate stiffness within the same sample over time reflects the progressive nature of pressure overload in hypertensive heart disease, overcoming the fundamental limitation of static afterload models that can only capture a single mechanical state (12). Furthermore, the non-invasive, magnetically actuated stiffening mechanism avoids the chemical perturbations associated with photo-crosslinkable hydrogels and the irreversibility of shape-memory elastomers, making MREs particularly suited for long-term biological studies (60, 67–69, 71). Finally, the combination of per-cell morphology, sarcomere organization, nuclear remodeling, NFAT3 signaling, fibroblast activation, and cell cycle analysis in a single platform provides an integrated, multi-readout view of mechanobiological remodeling in human myocardium.

Several limitations of this study should be acknowledged. First, the 2D monolayer format cannot fully replicate the 3D force transmission, cell-cell mechanical coupling, or anisotropic architecture of native myocardium or engineered heart tissues (108). Second, the degree of stiffening achievable with our current MRE formulation, while biologically effective, may not cover the full physiological range of pressure overload experienced *in vivo* (62). Third, our study focuses on a single HCM-causing mutation (*MYBPC3*fs), and it remains to be determined whether other sarcomeric mutations confer similar or distinct patterns of mechanosensitivity. Fourth, the opacity of MRE substrates makes real-time single-cell imaging (e.g. to assess how morphology evolves) technically challenging. Finally, 7-day stimulation window captures only early remodeling events; chronic mechanical stimulation over longer time may show additional genotype-specific maladaptations including metabolic reprogramming, fibrosis-like ECM remodeling, and electrophysiological dysfunction (109).

## Conclusion

In summary, this study establishes dynamically tunable MRE substrates as a physiologically relevant platform for modeling mechano-genetic interactions in HCM. By demonstrating that *MYBPC3*fs cardiomyocytes exhibit exaggerated hypertrophic remodeling, sarcomere disorganization, nuclear remodeling, dysregulated NFAT3 signaling, and aberrant cell cycle activity in response to mechanical stiffening, we provide direct evidence a *MYBPC3* loss-of-function genomic variant alter the cardiomyocyte mechanosensitivity. These findings advance our understanding of how genetic and mechanical stress may cooperate to initiate and amplify HCM pathology and open new path for dissecting disease-modifying pathways. Looking forward, this platform holds significant potential as a preclinical tool for testing therapeutic strategies aimed at reducing the cellular impacts of mechanical load, potentially by targeting calcineurin-NFAT and/or cell-cycle linked regulation in HCM.

## Methods

### MRE fabrication

MRE substrates were fabricated as described previously (68). Briefly, Sylgard 527 Dielectric Gel (Fisher Scientific; NC1208196; parts A and B were mixed 1:1 ratio) was combined with carbonyl iron particles (CIP; microspheres; Chemical Store; CAS 7439-89-6) and mixed by vigorous manual stirring for 10 min, followed by bath sonication (Branson 1510 Ultrasonic Cleaner) for 3 min to ensure homogeneous dispersion. The mixture was poured into an acrylic frame (**Fig. S1 F**), degassed for 10 minutes, and cured at 60°C for 24 hours. Substrates were prepared at 2.95 mm thickness with CIP loadings of 0, 33, 50, 66, 75, and 80 wt%; all cell experiments used 66 wt%. After curing, MREs were bonded to 22 mm × 22 mm glass coverslips using Sylgard 184 (Fisher Scientific; NC0162601; base and curing agent were mixed at 10:1 ratio) and post-cured for 4 hours at 60°C. Because cell studies require sterilization, a subset of MREs were autoclaved; baseline and magnetically induced stiffness were measured pre- and post-autoclave (68).

### Mechanical property characterization

#### Indentation testing

Spherical indentation was performed on an ElectroForce 3200 mechanical testing system (TA Instruments, New Castle, DE, USA) using a rigid 12 mm-diameter spherical indenter (radius *R*= 6 mm) mounted to the load axis and a 45 N load cell. Samples were placed on a custom 3D-printed poly(lactic acid) (PLA) stage that positioned a permanent magnet 2 mm beneath the substrate (**Fig. S1 A**). Because soft silicone gels adhere to indenters, substrate surfaces were conditioned with a thin, uniform film of a commercial hydrophobic coating (Rain-X, ITW Global Brands, Houston, TX, USA) and allowed to dry before testing, which kept the work of adhesion low and reproducible; adhesion was neglected in the analysis. Each test comprised six steps: (1) approach at 0.167 mm/s until a contact load of 0.02 N was registered (**Fig. S1 H, step 1**); (2) retraction to 0 N (**Fig. S1 H, step 2**); (3) a 25 s dwell (**Fig. S1 H, step 3**); (4) indentation at 0.167 mm/s to a depth of approximately 10% of substrate thickness (about 0.3 mm; **Fig. S1 H, step 4**); (5) a 100 s hold at constant depth during which force relaxation was recorded (**Fig. S1 H, step 5**); and (6) withdrawal until detachment. Force and displacement histories from steps (4) and (5) were analyzed as described next.

#### Viscoelastic analysis

Because the MRE is viscoelastic and beating cardiomyocytes load their substrate cyclically at approximately 1 Hz, we characterized each indentation site by its relaxation spectrum rather than by a single equilibrium modulus, and report the dynamic modulus at 1 Hz. The force history *F* (*t*) was related to the indentation depth *h* (*t*) through the Lee-Radok extension of Hertz contact to linear viscoelastic materials (110), which applies here because the contact area did not decrease during steps (4) and (5):

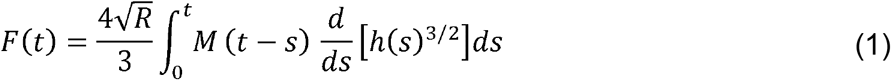

where *M* (*t*) is the reduced relaxation modulus of the substrate. The reduced relaxation modulus was represented by a generalized Maxwell model,

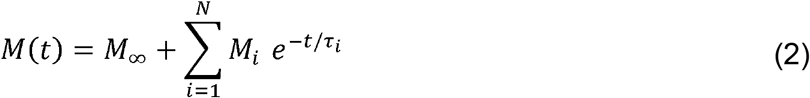

in which *M* _∞_ is the equilibrium reduced modulus and *M*_*i*_ is the amplitude of the Maxwell branch with relaxation time *τ* _*i*_. The relaxation times were prescribed as *N* = 100 values spaced logarithmically over 10^−2^ to 10^4^ s, so that Eq. (1) is linear in the unknown amplitudes *M* _∞_ and *M*_*i*_; these were estimated by least squares with Tikhonov regularization, with the regularization parameter chosen from the corner of the L-curve (111). Because the ramp lasted approximately 1.8 s and the hold 100 s, relaxation times between roughly 1 s and several hundred seconds are constrained directly by the data; shorter and longer relaxation times are constrained by the regularization and by the equilibrium term. Storage and loss reduced moduli at angular frequency follow analytically from the recovered spectrum:

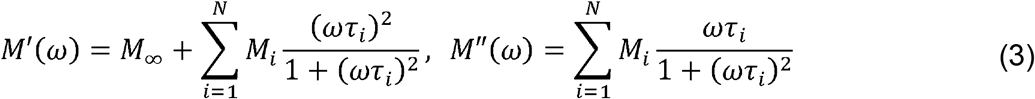

and the dynamic reduced modulus 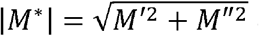 and loss tangent tan *δ* = *M* ^″^/ *M*′ were evaluated at *f* = *ω* /2 *π* = 1 Hz.

Because the substrate (thickness *h*_0_ = 2.95 mm) was bonded to a rigid coverslip and the contact radius 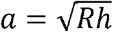 reached 1.3 mm, measured forces were corrected for finite thickness by dividing by the factor *f* = 1 + 1.133*χ* +1.283*χ*^2^ + 0.769*χ*^3^ + 0.0975*χ*^4^ for an incompressible layer bonded to a rigid substrate (112), evaluated at the full indentation depth (*χ* = *a*/*h*_0_ ≈ 0.45, *f* ≈ 1.85), at which the force was held throughout the relaxation phase that dominates the fit. The specimen was sufficiently thin that poroelastic effects were not expected to dominate (113). The reduced modulus is related to the Young’s modulus *E* and Poisson ratio *v* of the substrate by *M* =*E*/ (1− *v* ^2^), with the indenter treated as rigid; for an incompressible elastomer (*v* = 0.5), *E* = 3*M*/4 and the shear modulus is *G* = *M*/4. Sites at which the recovered spectrum yielded a negative loss tangent at 1 Hz (12 of 88), which can arise when the substrate has not fully recovered from handling or from a preceding indentation at an adjacent site, were excluded from the loss-tangent analysis. The regional contrast reported in Results was unchanged when these sites were retained.

#### Shear testing

Shear modulus was measured using a custom, quasi-static shear assay as described previously (68). Briefly, fishing wire (Ultra-Thin Diameter Superior Tensile T400 No. 8, Lazada, Philippines) was anchored with Sylgard 184 to the center of a 25 mm × 25 mm glass coverslip; the MRE was adhered to the opposite face. The free face of the MRE was attached to a 3D-printed PLA custom setup. On the opposite side of the setup, a hollow cylinder was made for placing permanent magnets, positioned 2 mm from the sample, same as the distance designed in the attachment for indentation testing. A binder clip of known mass was clamped to the fishing wire to apply a downward shear force. Images were captured immediately before and after loading with the smartphone camera aligned level with the sample plane to avoid parallax. Displacement of a particular point on the coverslip was quantified in ImageJ. Shear modulus *G* was approximated as:

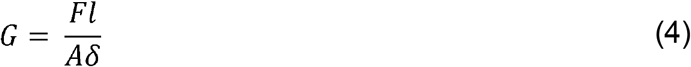

where *F* is applied force, *l* is the thickness of the specimen, *A* is the area over which shear force is distributed, and *δ* is displacement. Clip sizes were selected to produce measurable and non-detaching displacements (e.g., medium clip, 9.6 g, 94.1 mN at 0 mT; large clip, 27.9 g, 274 mN at 130 mT).

To quantify magnetically induced stiffening, the stiffness magnetorheological (SMR) effect was calculated as (77):

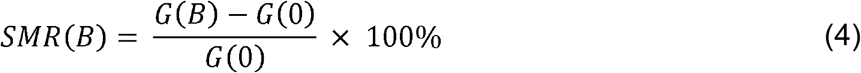

where *G* (*B*) is the shear modulus measured under the applied magnetic field *B* and *G* (0) G 0 is the shear modulus with no magnetic field applied.

### Stencil fabrication

Monolayer stencils were fabricated via Hydrogel-Assisted STereolithographic Elastomer (HASTE) replication (114). Briefly, initial CAD designs (Autodesk Inventor Professional) consisted of arrays of hollow circular through-holes (diameter: 6.4 mm) patterned into a 3 mm-thick base plate. Designs were printed in Clear Resin ™ on a Form 3 SLA 3D printer (Formlabs Inc., Somerville, MA). Printed resins were oxidized in an oxygen plasma chamber (Harrick Plasma, Ithaca, NY; 30 W, 90 s, 580–680 μTorr) to improve surface wettability and remove dust. Sylgard 184 was poured over the resin replica, clamped between two acrylic plates, and cured overnight at 60°C to yield ~1 mm-thick PDMS sheets containing circular through-holes. The cured PDMS sheets were detached from the replica, cut into individual circular stencils, and autoclaved. To minimize cell attachment to the stencils, sterilized stencils were incubated in 1% wt/vol Pluronic F127 for 1 hour at room temperature, followed by three 5 minutes washes in sterile dPBS.

### MRE surface modification

Stencils were briefly soaked in methanol, placed on sterilized MRE substrates, and kept in 60°C oven under septic condition overnight to promote physical adhesion via solvent evaporation (**Fig. S1 G**). Substrates were then coated with fibronectin (20 µg/mL in dPBS) for at least 2 hours to overnight, followed by three 5 minutes dPBS washes to remove unbound ECM.

### Cell culture

#### Cell viability (Alamar Blue)

To assess cytocompatibility of MRE substrates with different CIP wt% loadings, we performed an Alamar Blue viability assay using C2C12 myoblasts. First, sterilized MRE substrates were pre-incubated in C2C12 growth medium (high-glucose DMEM; Fisher Scientific; 12800082) supplemented with 10% FBS (Fisher Scientific; FB12999102), 1% penicillin-streptomycin (Fisher Scientific; 15140122), 1% GlutaMAX (Fisher Scientific; 35050061), and 1% non-essential amino acids (Fisher Scientific; 11140050) for 72 hours at 37°C. The conditioned medium from each substrate group was collected for testing. In parallel, C2C12 cells were seeded at 40,000 cells/cm^2^ in a 96-well plate and cultured for 24 hours. At this point, their culture medium was incubated with 10% (v/v) Alamar Blue reagent (Fisher Scientific; Y00-100) for 2 hours before plate reader measurement, which was used as the baseline fluorescence signal. Next, the conditioned media collected from MRE substrates was applied to C2C12 cells in 96-well plates and cultured for 24 hours. Media from these wells was then incubated with 10% (v/v) Alamar Blue reagent for 2 hours and measured with the plate reader. Normalized fluorescence intensity was calculated as the ratio of fluorescence captured in conditioned media over fluorescence captured in baseline.

#### Human primary ventricular cardiac fibroblasts

Human primary ventricular cardiac fibroblasts (Lonza; CC-2904) were expanded in fibroblast growth medium (FBM; Fisher Scientific; NC0089421) supplemented with 10% FBS, 0.1% insulin, 0.1% hFGF-B, and 0.1% GA-1000 to ~90% confluency in T25 flasks, dissociated with 0.25% trypsin–EDTA (Fisher Scientific; 25200114) for 5 minutes, quenched with FBM, and replated on MREs at 10,000 or 25,000 cells/cm^2^ in FBM.

#### Human induced pluripotent stem cells derived cardiomyocytes

Isogenic control iPSC derived from a healthy male with no family history of heart disease (Coriell repository # GM25256) were previously edited to express a single genomic copy of the genetic Ca^2+^ indicator GCaMP6f (115), to optically measure cardiomyocytes’ calcium transients. A patient-specific *MYBPC3* frameshift mutation identified to be pathogenic, (*MYBPC3* <u>G1206</u>fs, *MYBPC3*fs (74)), was engineered into the Control-GCaMP6 background (75).

Isogenic control and *MYBPC3*fs hiPSCs were cultured in StemFlex media (Fisher Scientific; A3349401) on Geltrex-coated plates (18.75 μg/cm^2^; Fisher Scientific; A1413302). For passaging and differentiation, cells were dissociated to single cells with Gentle Dissociation Reagent (PBS with 1.8 g/L NaCl and 0.5 mM EDTA; (116)) and replated with 10 μM Y-27632 (Biogems; 1293823).

Cardiomyocyte differentiation was induced by small-molecule Wnt modulation (54, 117). Human iPSCs were seeded at 40,000 cells/cm^2^ on Geltrex (37.5 μg/cm^2^) and expanded 3 days to 90% confluency. On day 0, media was changed to RPMI1640 (Life Technologies; 11875093) supplemented with 150 μg/ml ascorbic acid (Fisher Scientific; NC0602549) and 2% B27 without insulin (Life Technologies; A1895601) (RPMI-I) containing 6 μM CHIR99021 (Biogems; 2520691) (differentiation day 0). After 2 days, media was changed to RPMI-I containing 5 μM IWP2 (Biogems; 6866167). After 2 additional days, media changed to RPMI-I. At day 6 of differentiation, media was changed to RPMI1640 supplemented with 2% B27 (Life Technologies; 17504044) (RPMI-C). Thereafter, culture was fed every 2 days with RPMI-C. Spontaneous beating typically appeared by day 8. On day 14, monolayers were gently dissociated with 10X TrypLE Select (Gibco; A1217701) and replated on Geltrex-coated plastic (37.5 μg/cm^2^) at 300,000 cells/cm^2^ in RPMI-C supplemented with 20% FBS and 10 μM Y-27632 for 24 h. On day 15, metabolic selection was initiated with glucose-free RPMI (Millipore Sigma; R1383) supplemented with 4 mM lactate (Fisher Scientific; AAL1450014), 1% NEAA, and 1% GlutaMAX. Medium was refreshed on day 18. From days 21–24, cultures were gradually transitioned back to RPMI-C. Cardiomyocytes purities were quantified using flow cytometry analysis of cardiac troponin T positive cells. Purified hiPS-CMs (days 24–26) were then seeded on MRE substrates at 50,000 cells/cm^2^ in RPMI-C supplemented with 10% FBS and 10 μM Y-27632.

#### Mechanical stimulation

Five days after monolayer formation, neodymium grade N52 magnets (K&J Magnetics; BX084TP-N52) were positioned to chronically apply static magnetic fields to subset of samples (human primary ventricular cardiac fibroblast or hiPS-CM cultures on 66 wt% MREs) to apply static magnetic field (~64 mT at the center of the substrates; **Fig. S1B**) for 7 days. Media was changed every 3 days.

#### Chemical stimulation

To induce fibroblast-to-myofibroblast transition, primary cardiac fibroblast cultures on MREs were treated with 10 ng/ml TGF-β (PeproTech; AF-100-36E) for 7 days. Media was changed every 3 days. Fresh TGF-β was added with each new media addition.

### Immunostaining

Monolayers were incubated in 100 mM KCl until cessation of spontaneous beating and then fixed with 4% paraformaldehyde at 4°C for 1 hour and washed in dPBS three times each time for 5 minutes. Primary antibodies included sarcomeric α-actinin (Millipore Sigma; A7811), pan-cadherin (Cell Signaling; 5332), NFAT3 (Fisher Scientific; A302-769A), and Ki67 (Fisher Scientific; 41-5698-82) for hiPS-CMs; and phalloidin (Fisher Scientific; A22287) and α-SMA (Abcam; ab124964) for fibroblast/myofibroblast markers. Cell nuclei were stained with Hoechst 33342. Monolayer stained were kept in dPBS and imaged using a Fluoview FB1000 confocal microscope (Olympus, Tokyo, Japan). To enable quantitative comparison of NFAT3, Ki67, and α-SMA expression across different conditions, imaging parameters acquisition were held constant as table 1:

**Table 1.** Confocal imaging acquisition parameters used for quantitative analysis of NFAT3, Ki67, and α-SMA immunofluorescence. Imaging settings were held constant across all experimental conditions to enable quantitative comparison of fluorescence intensity.

| Antibody | Laser wavelength (nm) | Transmissivity (%) | PMT (V) | Gain | Offset (%) | Pixels | Objective |
| --- | --- | --- | --- | --- | --- | --- | --- |
| NFAT3 | 488 | 2.8 | 600 | 1 | 8 | 2048×2048 | 20X |
| Ki67 | 488 | 3 | 510 | 1 | 8 | 2048×2048 | 20X |
| α-SMA | 488 | 2 | 500 | 1 | 8 | 1042×1042 | 10X |

### Structural analysis

Cell boundary segmentations were performed using the CellPose (Cyto2 model) (118) and sarcomere analysis was performed using SarAsM (84), and structural analysis codes modified from the original form described in (108) are available here: [Link withheld for review].

### Quantification and statistical analysis

GraphPad 9.3.1 was used for statistics and data visualization. Data are presented with mean ± standard deviation (*SD*) and are pooled from 2D monolayers from at least 3 independent differentiation batches. For quantitative analysis, data were first assessed for normality using Normality and Lognormality tests. If datasets were normally distributed, comparisons between two groups were performed using a parametric Student’s t-test, and comparisons among multiple groups were analyzed using parametric one-way ANOVA followed by Holm–Šidák post hoc test for multiple comparisons. When two independent variables were compared, data were analyzed using two-way ANOVA followed by Holm–Šidák post hoc test. If datasets were not normally distributed, a non-parametric Mann–Whitney U test was used for two-group comparisons, and one-way ANOVA, a Kruskal–Wallis test, followed by Dunn’s multiple comparison was applied for multiple group comparisons. For genotype × magnetization experiments, the four groups were compared using one-way ANOVA with Holm–Šidák post hoc test (normally distributed data) or a Kruskal–Wallis test with Dunn’s multiple comparisons test (non-normally distributed data). The spatial trend in stiffness was assessed with a one-sided Page’s L test (Supplementary Section 1.6). All statistical analyses were performed as two-tailed tests unless otherwise specified.

## Acknowledgments

We thank the Genome Engineering & Stem Cell Center (GESC@MGI) at Washington University in St. Louis for cell line engineering services. We also thank members of the Huebsch’s laboratory for input, support, and guidance during the completion of this work and preparation of the manuscript. We would also like to thank Dr. Michael Vahey of the Washington University in Saint Louis McKelvey School of Engineering for his support and guidance on laser scanning confocal microscopy, Dr. Gretchen Meyer of the Washington University in Saint Louis School of Medicine for generously providing C2C12 cells, and Barbra Semar of the Washington University in Saint Louis McKelvey School of Engineering for her support and guidance with mechanical testing.

## Author contributions

Conceptualization: YKGK, YS, SC, GMG, NH

Methodology: YKGK, YS, GM, FL, GMG, NH

Investigation: YKGK, YS, LH, FL, NDD, HJ

Formal analysis: YKGK, YS, BB

Software: YKGK, BB, GM, NH

Visualization: YKGK, NH

Supervision: GMG, NH

Writing—original draft: YKGK, SC, GMG, NH

Writing—review and editing: YKGK, SC, GMG, NH

