## Supplementary Material for "Tunable Magnetorheological Elastomer Substrates for Studying Mechano-Genetic Interactions in Hypertrophic Cardiomyopathy"

Whitaker Hall 300E

1 Brookings Drive

Saint Louis, MO, 63130

1.1 Cell Segmentation with CellPose (Cyto2) model:

Nuclei and α-actinin stained images were paired and organized prior to segmentation, ensuring that only correctly matched image sets were processed. Images were converted to float32, normalized to a [0, 1] range, and stacked into two-channel arrays with α-actinin assigned as the first channel to ensure accurate interpretation by CellPose. Segmentation was performed using the cyto2 model in 2D mode with an initial diameter of 100 pixels. Resulting masks were used to extract cellular morphology features including:

1) area (μm^2^)
2) elongation = $\frac{MajorAxisLength}{MinorAxisLength}$
3) eccentricity = $\sqrt{1-\left( \frac{MinorAxisLength}{MajorAxisLength} \right)^{2}}$
4) circularity = $\frac{4\pi\times area}{Perimeter^{2}}$

Measurements were exported to a single CSV file, and false-color overlays were generated with region IDs to enable visual quality control. Two strategies were applied to optimize segmentation. First, the CellPose cyto2 model was used directly while adjusting the estimated cell diameter to improve agreement between predicted and observed a-actinin-positive cell boundaries. Although this approach improved segmentation in some images, defining cell boundaries solely by a-actinin-positive staining remained challenging because the signal contains both cortical cell-border information and intracellular sarcomeric structures (Pre-Training; **Fig. S2**).

Second, the Cellpose Cyto2 model was fine-tuned using manually annotated membrane-stained images to improve segmentation of a-actinin-positive cells. Briefly, fluorescence images stained with a membrane-associated protein (Pan-Cadherin) were manually annotated to generate reference cell-boundary masks. Each membrane image was paired with its corresponding manual segmentation mask and used to fine-tune the pretrained Cyto2 model. During training, image patches containing cells were extracted, normalized, and augmented to improve model robustness, while the model weights were iteratively updated by minimizing the difference between predicted segmentations and the manually annotated reference masks. Following optimization on membrane-stained images, the fine-tuned model was applied to paired nuclei and a-actinin datasets. Because the model had learned cell-boundary features from the membrane annotations, it was able to more accurately delineate cardiomyocyte boundaries in a-actinin-positive cells despite the presence of prominent intracellular sarcomeric structures. Compared with the pretrained Cyto2 model, the fine-tuned model produced more accurate and consistent segmentation of a-actinin-positive cells (Post-Training; **Fig. S2**).

1.2 Calculation for a-SMA expression

To calculate α-SMA expression in cardiac fibroblasts (cFBs), individual cells were first segmented using CellPose cyto2 model. In this process, the α-SMA-positive staining was used to identify individual cell regions. Cell boundaries were identified directly from α-SMA immunofluorescence images, and the estimated cell diameter was set to 100 pixels to optimize agreement between the predicted segmentation masks and observed average size of α-SMA-positive cells. The resulting segmentation masks were used to define individual cell boundaries for subsequent quantification of α-SMA mean fluorescence intensity per cell.

1.3 Nucleus Segmentation with CellPose Nucleus Model

Nuclei were segmented from Hoechst-stained images using the Cellpose nucleus model. Prior to segmentation, nuclei images were preprocessed by percentile-based intensity normalization (1–99.8%), followed by contrast-limited adaptive histogram equalization (CLAHE) and light Gaussian smoothing to improve nuclear detection across fields with variable staining intensity. CLAHE was applied to enhance local image contrast, improving detection of dim nuclei without substantially increasing background intensity, while Gaussian smoothing reduced high-frequency image noise and improved segmentation consistency across fields with variable staining intensity. This preprocessing pipeline was applied exclusively for nuclear segmentation and did not alter the raw fluorescence intensities used for downstream protein quantification. The resulting nuclear masks were used to extract nuclear morphology features. Although preprocessing improved segmentation accuracy, nuclei within regions of strong signal saturation or poor image contrast were not reliably detected and were excluded from subsequent analysis.

1.4 Sarcomere using SarcAsM

Sarcomere organization was quantified using the SarcAsM pipeline (1) on paired Hoechst and a-actinin fluorescence images. Cardiomyocytes were first segmented using the Cellpose Cyto2 model with Hoechst and a-actinin (as in sections 1.1 and 1.3). Prior to analysis, the a-actinin channel was enhanced using an Unsharp Mask followed by Gaussian high-pass filtering to improve local contrast and emphasize Z-disc structures. SarcAsM then identified individual Z-discs and generated vectors connecting neighboring Z-discs, from which local sarcomere length (Z-disc spacing) and orientation were calculated. Z-discs were subsequently grouped into connected components within each cell, and each component was classified as organized or disorganized based on local vector statistics, including sarcomere length, orientation consistency, and minimum vector and repeat requirements. In this study, components were classified as organized if they showed a mean sarcomere length between 0.8-2.2 μm, an orientation standard deviation $\leq$20°, at least three vectors per component, and a minimum of three sarcomere repeats per vector. Components failing to satisfy these criteria were classified as disorganized. To avoid bias from image artifacts, a hotspot mask identifying saturated and high-intensity pixels was generated, and components with $\geq$10% hotspot overlap were excluded from analysis. The pipeline exported component-level, cell-level, and image-level metrics together with quality-control overlays. Because hiPSC-CMs were cultured as a 2D monolayer without global alignment, subsequent analyses focused on the Component model, which quantifies short-range Z-disc organization within individual cells.

1.5 NFAT3 subcellular localization analysis

NFAT3 subcellular localization was quantified from three-channel fluorescence images: Hoechst (nuclei), sarcomeric α-actinin (cell boundaries), and NFAT3 (signal of interest). Nuclear segmentation was performed as described in section 1.3, and cell boundary segmentation was performed as described in section 1.1.

Nuclear and cell masks were merged by assigning each segmented nucleus to the cell with which it shared the greatest pixel overlap. Nuclear pixels were subsequently relabeled with the corresponding cell ID, which enabled identification of the nuclear and cytoplasmic compartments for each cardiomyocyte. The cytoplasmic compartment was defined as the segmented cell area excluding the nuclear region. Cells without an assigned nucleus were excluded from subsequent analysis.

NFAT3 fluorescence intensity was quantified from the raw NFAT3 channel without any normalization. Mean and total NFAT3 fluorescence intensities were recorded separately for the nuclear and cytoplasmic compartments of each cell; for display (**Fig. 4D, E**), per cell MFI values were normalized to the mean of the unmagnetized isogenic control. Nuclear translocation was calculated as the nuclear-to-cytoplasmic (nuc/cyto) NFAT3 fluorescence intensity ratio:

$$NFAT3 nuclear translocation= \frac{raw NFAT3 intensity in nucleus}{raw NFAT3 intensity in cytoplasm}$$

Segmentation quality was verified by overlaying cell boundaries and nuclear boundaries on the NFAT3 grayscale channel to confirm correct compartment assignment prior to quantification.

1.6 Frequency-domain characterization at 1 Hz

Indentation force–displacement data were analyzed using a generalized Maxwell model coupled with the Hertz–Lee–Radok contact formulation,

$$F(t) = (4\surd R/3) \int₀ᵗ M(t - s) (d/ds)[h(s)^{3}ᐟ^{2}] ds,$$

where $R$ is the indenter radius, $h(t)$ is indentation depth, and $M(t)$ is the reduced relaxation modulus.

$$M(t) = M\infty+ \sumᵢ₌₁ᴺ Mᵢ e^(-t/\tauᵢ).$$

Here, $M\infty$ is the equilibrium reduced modulus, and $M_{i}$ is the reduced-modulus amplitude of the Maxwell branch with relaxation time $\tau_{i}$. The discrete relaxation spectrum consists of these amplitudes at N = 100 logarithmically spaced relaxation times spanning 10^-2^–10^4^ s.

Storage and loss reduced moduli were calculated analytically from the recovered generalized Maxwell spectra at$f = 1 Hz$using

$$M'(\omega) = M\infty+ \sumᵢ₌₁ᴺ Mᵢ (\omega\tauᵢ)^{2}/[1 + (\omega\tauᵢ)^{2}]$$

$$M''(\omega) = \sumᵢ₌₁ᴺ Mᵢ \omega\tauᵢ/[1 + (\omega\tauᵢ)^{2}].$$

The dynamic reduced modulus and loss tangent were then obtained as

$$|M*(\omega)| = \surd([M'(\omega)]^{2} + [M''(\omega)]^{2}) and tan(\delta) = M''(\omega)/M'(\omega).$$

The spatial trend in dynamic reduced modulus at 1 Hz was assessed using a one-sided Page test across 11 positions (0–10 mm), with four independent specimens as blocks. Two measurement series were averaged at each specimen–position. Within each specimen, modulus values were ranked from 1 (lowest) to 11 (highest), and Page’s L was calculated using increasing position weights from the center to the edge. The unadjusted p-value was reported, with significance defined as *p* < 0.05.

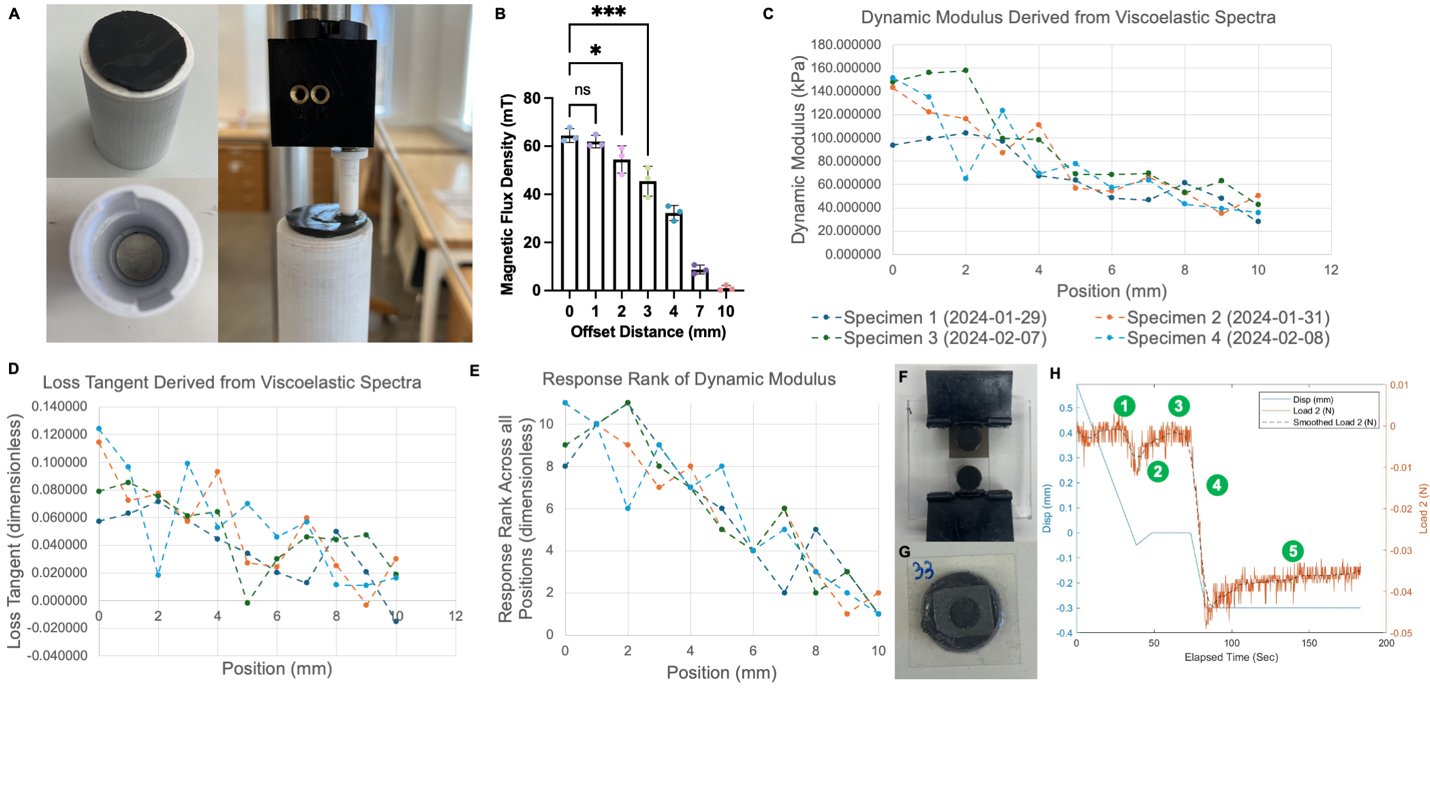

**Figure S1. Spatial characterization of magnetically induced stiffening across the MRE surface.** **(A)** Custom non-magnetic indentation setup used to measure local MRE stiffness at defined lateral offsets from the center MRE (magnetized condition). Insets show the indenter tip, MRE, and magnet configuration. **(B)** Magnetic flux density measured as a function of lateral distance from the center of MRE, demonstrating a progressive decline in magnetic field strength with increasing offset. Statistical significance was determined by one-way ANOVA with Dunnett multiple comparisons test; * *p* < 0.05, ** *p* < 0.001; ns, not significant. **(C)** Dynamic reduced modulus at 1 Hz versus distance from the specimen center (0–10 mm) for four independent specimens. Each point averages two measurement series. **(D)** Loss tangent at 1 Hz versus distance from the specimen center, for the same four independent specimens; each point averages two measurement series. **(E)** Corresponding within-specimen ranks, from 1 (lowest modulus) to 11 (highest). The decreasing spatial trend was assessed using a one-sided Page’s test (panels C-E; *L* = 1177, unadjusted $=$ $\times$ 9.09 ￼ 10^-16^). **(F)** Representative MRE. **(G)** Representative MRE substrate with the stencil positioned on top within the region of uniform magnetic actuation (6.4 mm diameter). **(H)** Representative indentation trace illustrating the testing protocol used to calculate elastic modulus. The blue trace represents indenter displacement and the orange trace represents measured load. The numbered steps indicate (1) probe approach and contact detection, (2) probe retraction to remove preload, (3) 25-s dwell at zero load, (4) indentation to the target depth, and (5) force-relaxation hold used viscoelastic spectrum fitting.

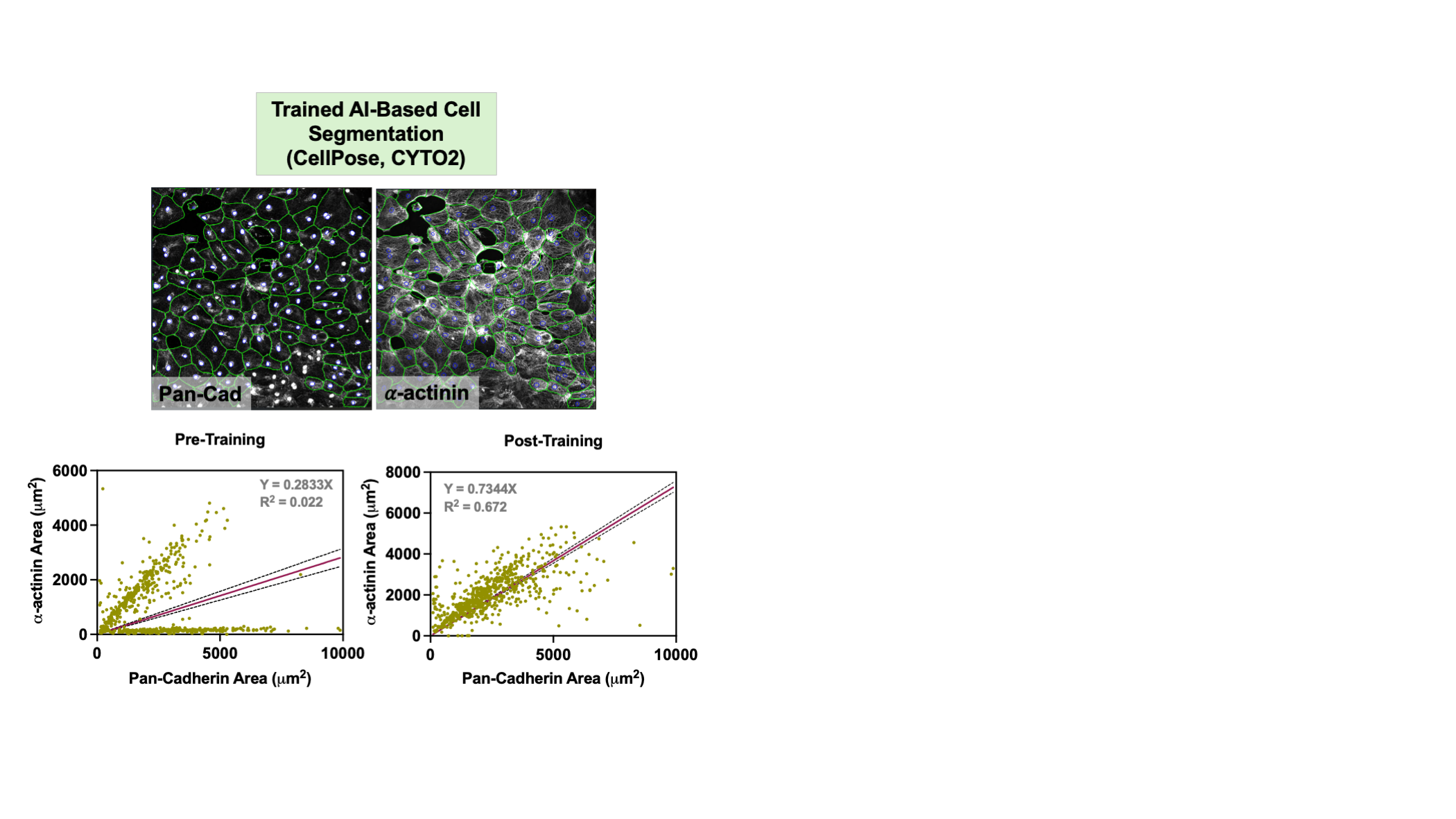

**Figure S2. AI-based cell segmentation using cortical 𝛼-actinin staining to detect cardiomyocyte boundaries.** Representative images of hiPSC-CM monolayers showing CellPose (Cyto2 model) segmentation outputs (green outlines) using Pan-Cadherin and 𝛼-actinin as cell boundary markers, with Hoechst-stained nuclei (segmented in blue). Scatter plots show the correlation between Pan-Cadherin-defined cell area and 𝛼-actinin-defined cell area before (Pre-Training, left) and after (Post-Training, right) model training. Prior to training, segmentation using 𝛼-actinin yielded poor agreement with Pan-Cadherin-defined cell area (R^2^ = 0.022), indicating inaccurate boundary detection. After fine-tuning the Cyto2 model on cortical 𝛼-actinin staining, segmentation accuracy improved substantially (R^2^ = 0.672), demonstrating that the trained model reliably identifies cardiomyocyte boundaries using 𝛼-actinin alone without reliance on membrane staining.

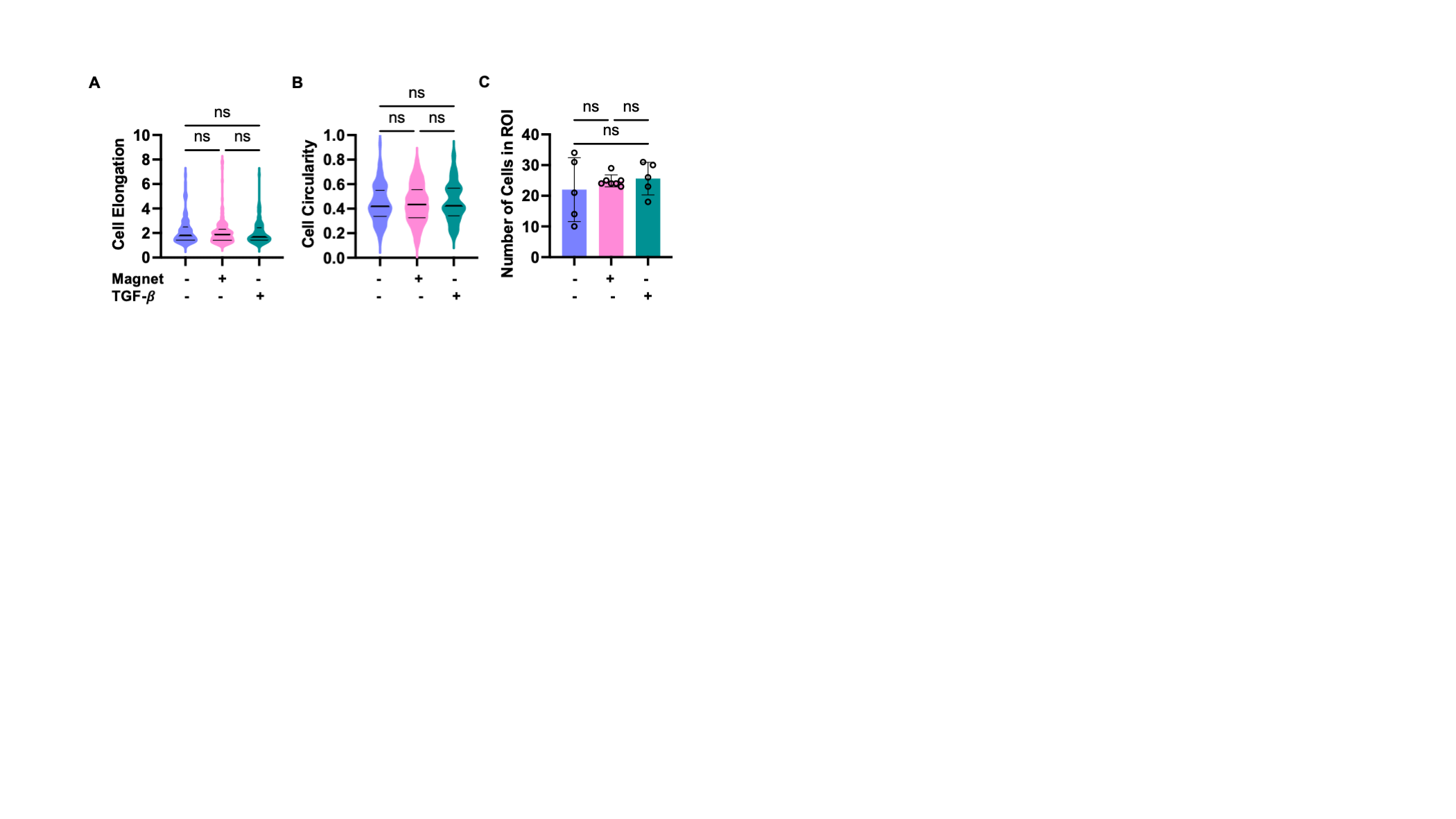

**Figure S3. Mechanical and chemical stimulation do not significantly alter cardiac fibroblast elongation, circularity, or cell density.** Quantification of **(A)** cell elongation, **(B)** cell circularity, and **(C)** number of cells per region of interest (ROI) across CTRL, Magnet, and TGF-𝛽 conditions. No significant differences were observed in any morphological parameter or cell proliferation among the three groups. Data in **(A, B)** are presented as violin plots, n represent individual cells; data in **(C)** are presented as mean $\pm$*SD,* n represent individual ROI. *P* values by one-way ANOVA, non-parametric, Kruskal-Wallis test with Dunn's multiple comparisons; ns = not significant.

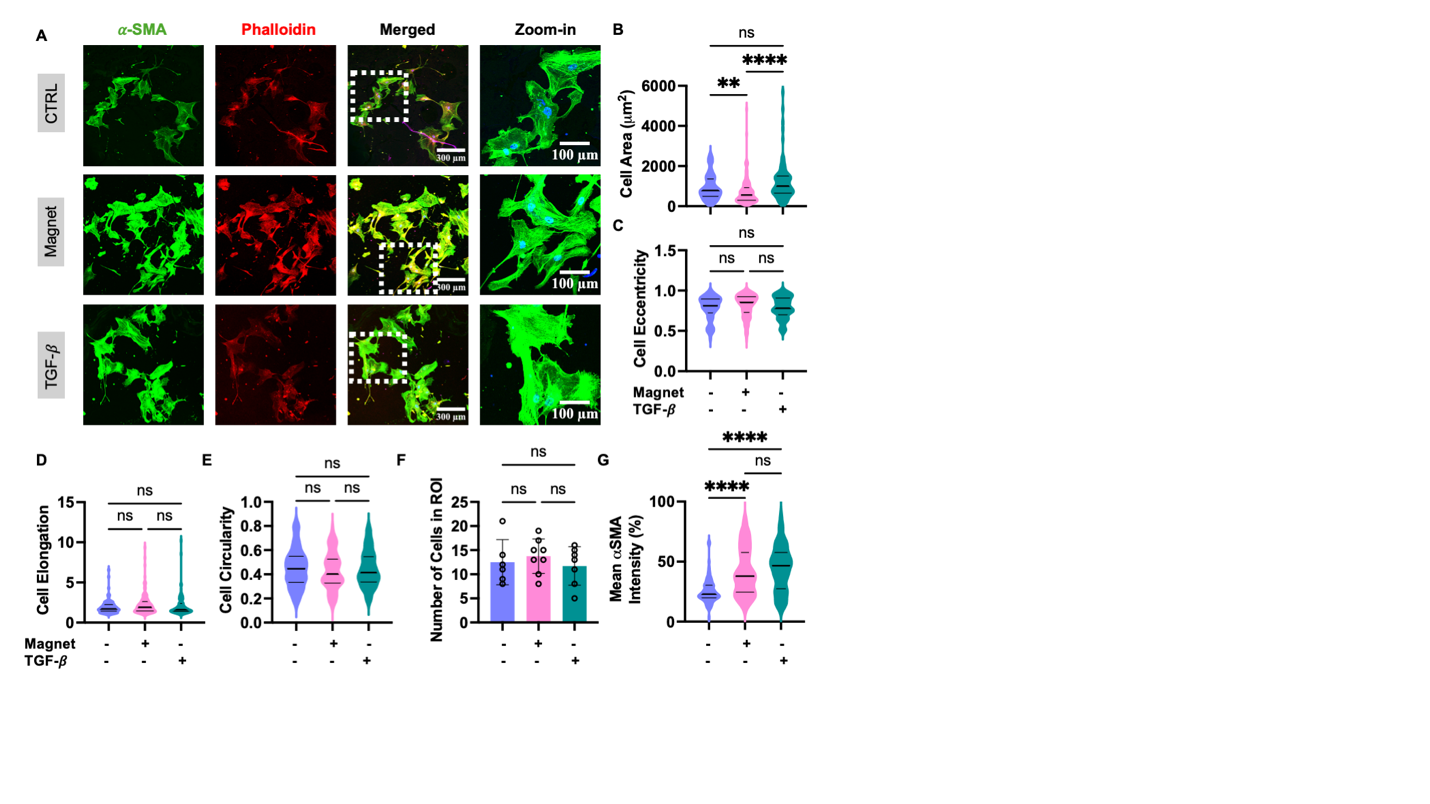

**Figure S4. Mechanical and chemical stimulation differentially regulate cardiac fibroblast morphology and activation at a seeding density of 10,000 cells/cm^2^. (A)** Representative immunofluorescence images of primary cardiac fibroblasts under three conditions; control (CTRL), mechanical stimulation (Magnet), and chemical stimulation (TGF-β) — stained for 𝛼-SMA (green), Phalloidin (red), and Hoechst (blue), shown as individual channels, merged, and zoomed-in views. Scale bars: 300 𝜇m (merged) and 100 𝜇m (zoom-in). Quantification of **(B)** cell area, **(C)** cell eccentricity, **(D)** cell elongation, **(E)** cell circularity, **(F)** number of cells per ROI, and **(G)** mean 𝛼-SMA intensity (%) across CTRL, Magnet, and TGF-𝛽 conditions. TGF-𝛽 treatment significantly increased 𝛼-SMA expression compared to CTRL, and mechanical stimulation produced an increase in 𝛼-SMA expression comparable to TGF-𝛽. No significant differences were observed in cell eccentricity, elongation, circularity, or cell density across any condition. Data in (**B–E**) and (G) are presented as violin plots, n represent individual cells; data in (**F**) are presented as mean $\pm$ *SD,* n represent individual ROI. *P* values by one-way ANOVA, non-parametric, Kruskal-Wallis test with Dunn's multiple comparisons; **** *p* < 0.0001, ** *p* < 0.01, ns = not significant.

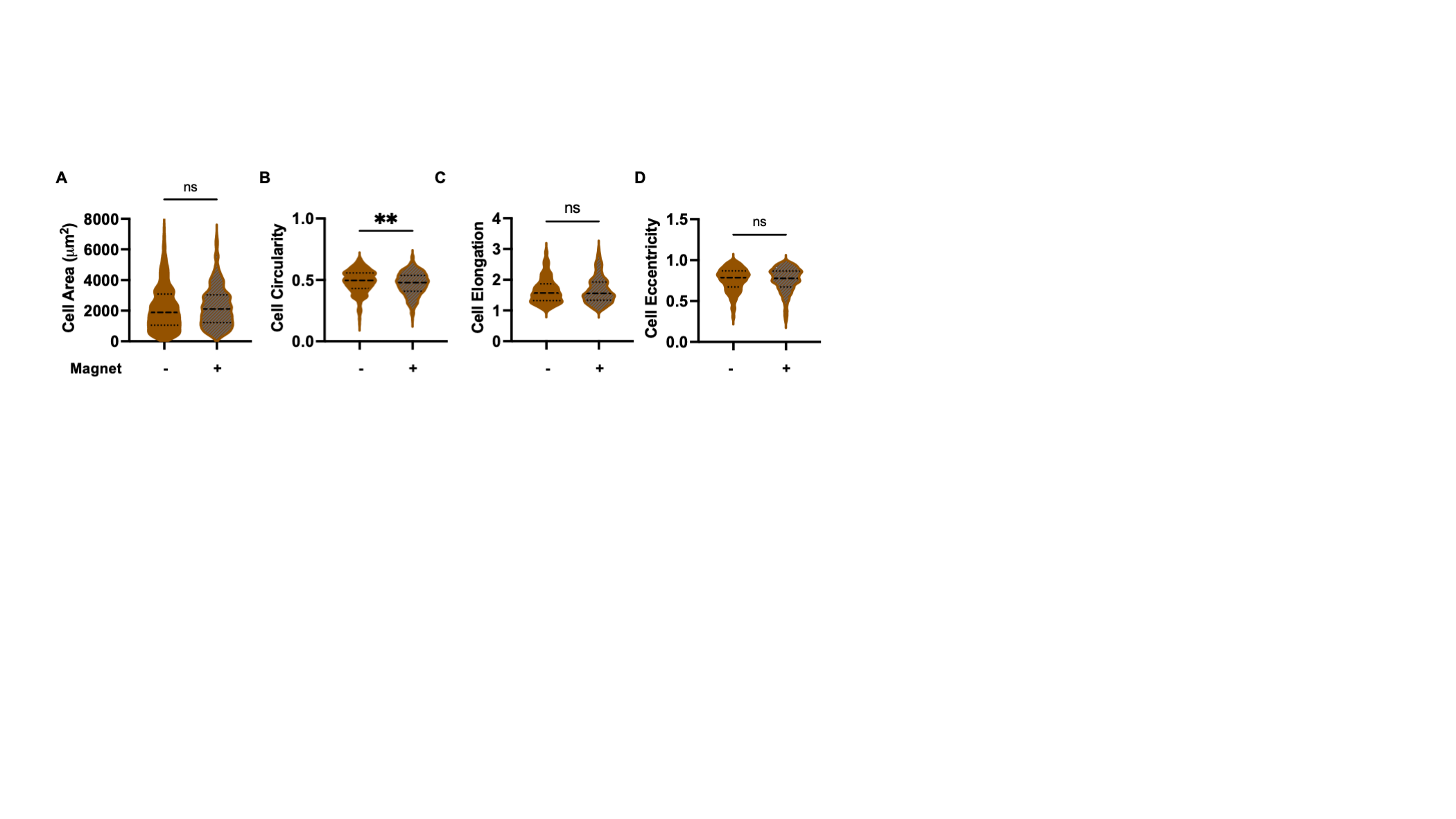

**Figure S5. Mechanical stimulation induces modest morphological changes in isogenic control (WT) hiPSC-CMs.** Quantification of **(A)** cell area, **(B)** cell circularity, **(C)** cell elongation, and **(D)** cell eccentricity in WT hiPSC-CMs under unmagnetized (−) and magnetized (+) conditions. Magnetization produced a modest but significant decrease in cell circularity, while cell area, elongation, and eccentricity were not significantly altered. Data are presented as violin plots, n represent individual cells; *P* values by unpaired t-test, non-parametric, Mann-Whitney test; ** *p* < 0.01, ns = not significant.

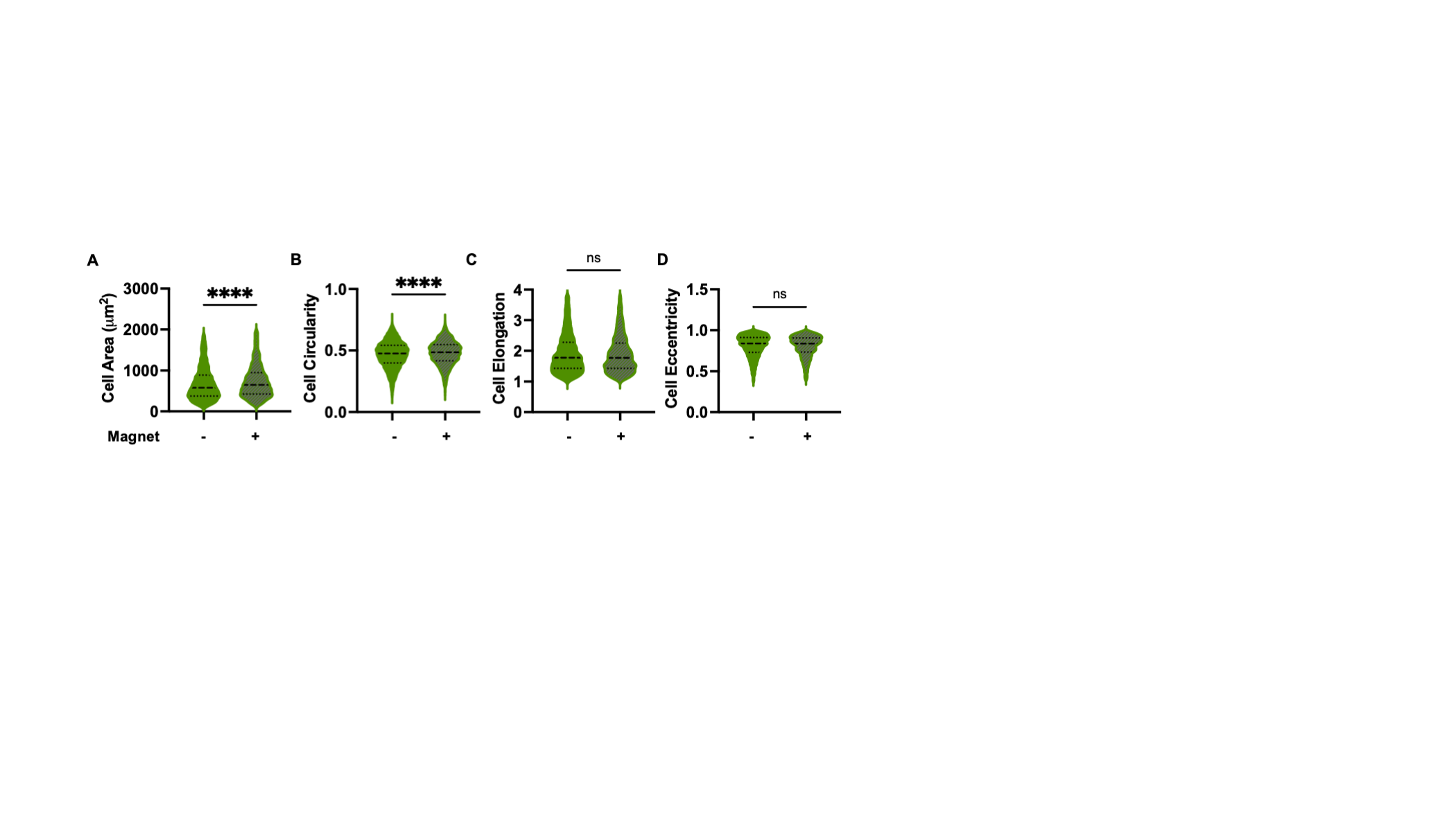

**Figure S6. Mechanical stimulation induces hypertrophic morphological remodeling in *MYBPC3fs* hiPSC-CMs.** Quantification of **(A)** cell area, **(B)** cell circularity, **(C)** cell elongation, and **(D)** cell eccentricity in *MYBPC3fs* hiPSC-CMs under unmagnetized (−) and magnetized (+) conditions. Magnetization significantly increased both cell area and cell circularity in *MYBPC3fs* cardiomyocytes, indicating a more isotropic, hypertrophic morphology, while cell elongation and eccentricity were not significantly altered. Data are presented as violin plots, n represent individual cells; *P* values by unpaired t-test, non-parametric, Mann-Whitney test; **** *p* < 0.0001, ns = not significant.

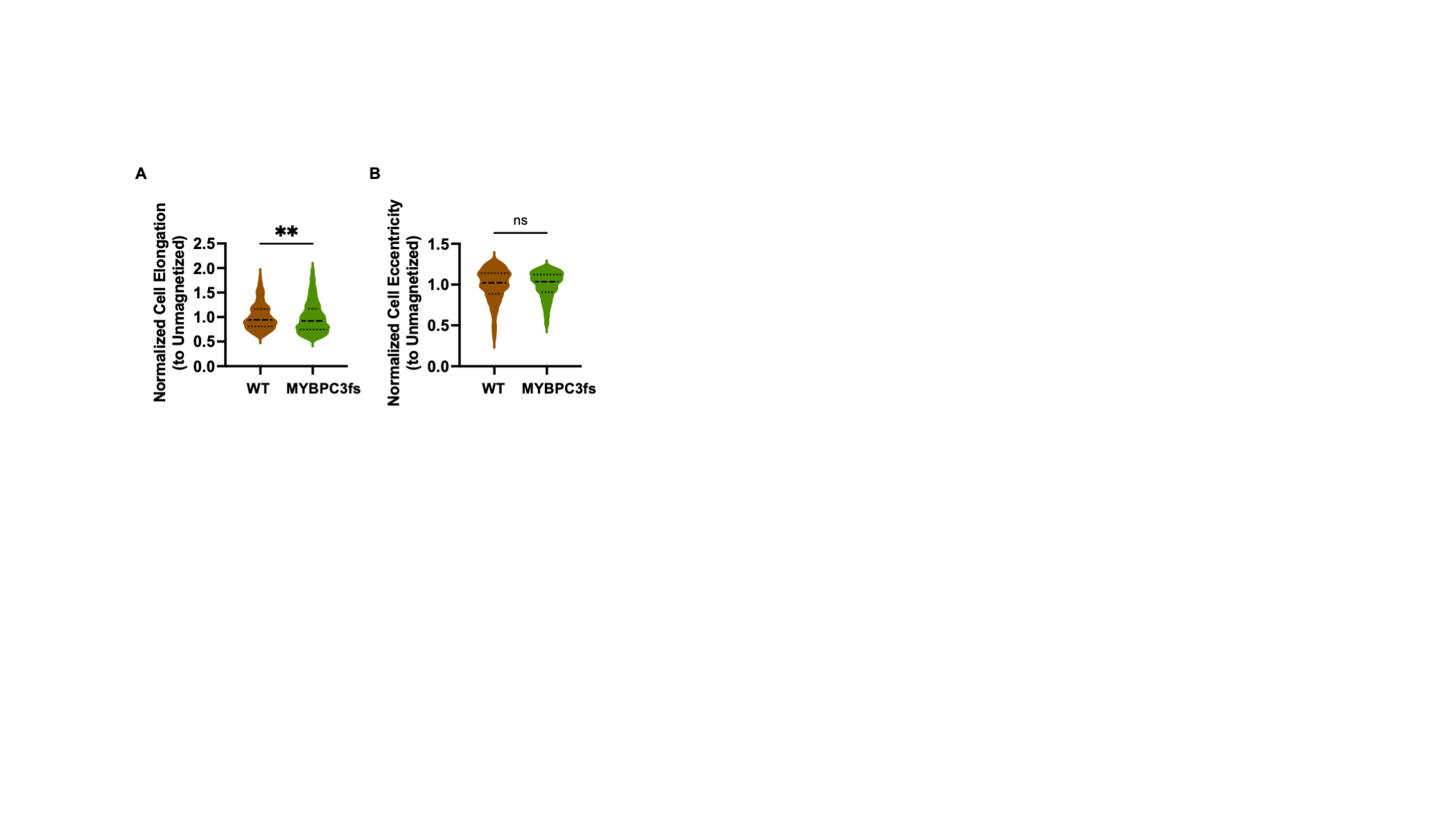

**Figure S7. *MYBPC3fs* cardiomyocytes show greater magnetization-induced changes in cell elongation compared to isogenic WT controls.** Quantification of **(A)** normalized cell elongation and **(B)** normalized cell eccentricity relative to respective unmagnetized controls for WT and *MYBPC3fs* hiPSC-CMs following magnetization. *MYBPC3fs* cardiomyocytes showed a significantly greater reduction in cell elongation upon magnetization compared to WT controls, while normalized cell eccentricity did not differ significantly between genotypes. Data are presented as violin plots, n represent individual cells; *P* values by unpaired t-test, non-parametric, Mann-Whitney test; ** *p* < 0.01, ns = not significant.

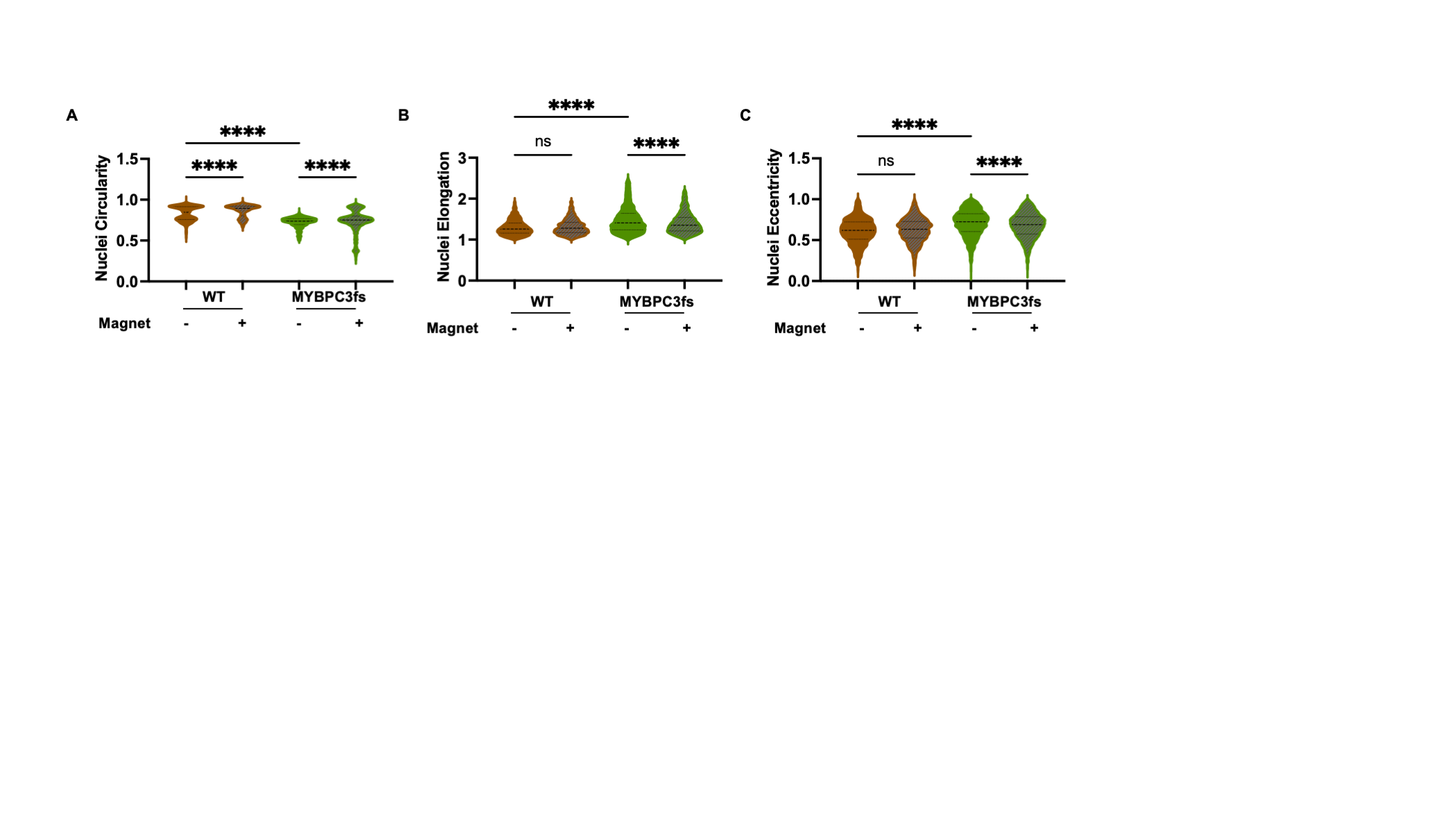

**Figure S8. Mechanical stimulation alters nuclear morphology in a genotype-dependent manner in hiPSC-CMs.** Quantification of **(A)** nuclear circularity, **(B)** nuclear elongation, and **(C)** nuclear eccentricity across WT and *MYBPC3fs* hiPSC-CMs under unmagnetized (−) and magnetized (+) conditions. In WT cardiomyocytes, magnetization significantly increased nuclear circularity, while nuclear eccentricity and elongation were unchanged. In *MYBPC3fs* cardiomyocytes, magnetization significantly reduced nuclear eccentricity and elongation while increasing nuclei circularity, indicating a shift toward a rounder nuclear morphology upon mechanical stimulation. Data are presented as violin plots, n represent individual nuclei; *P* values by one-way ANOVA, non-parametric, Kruskal-Wallis test with Dunn's multiple comparisons; **** *p* < 0.0001, ns = not significant.

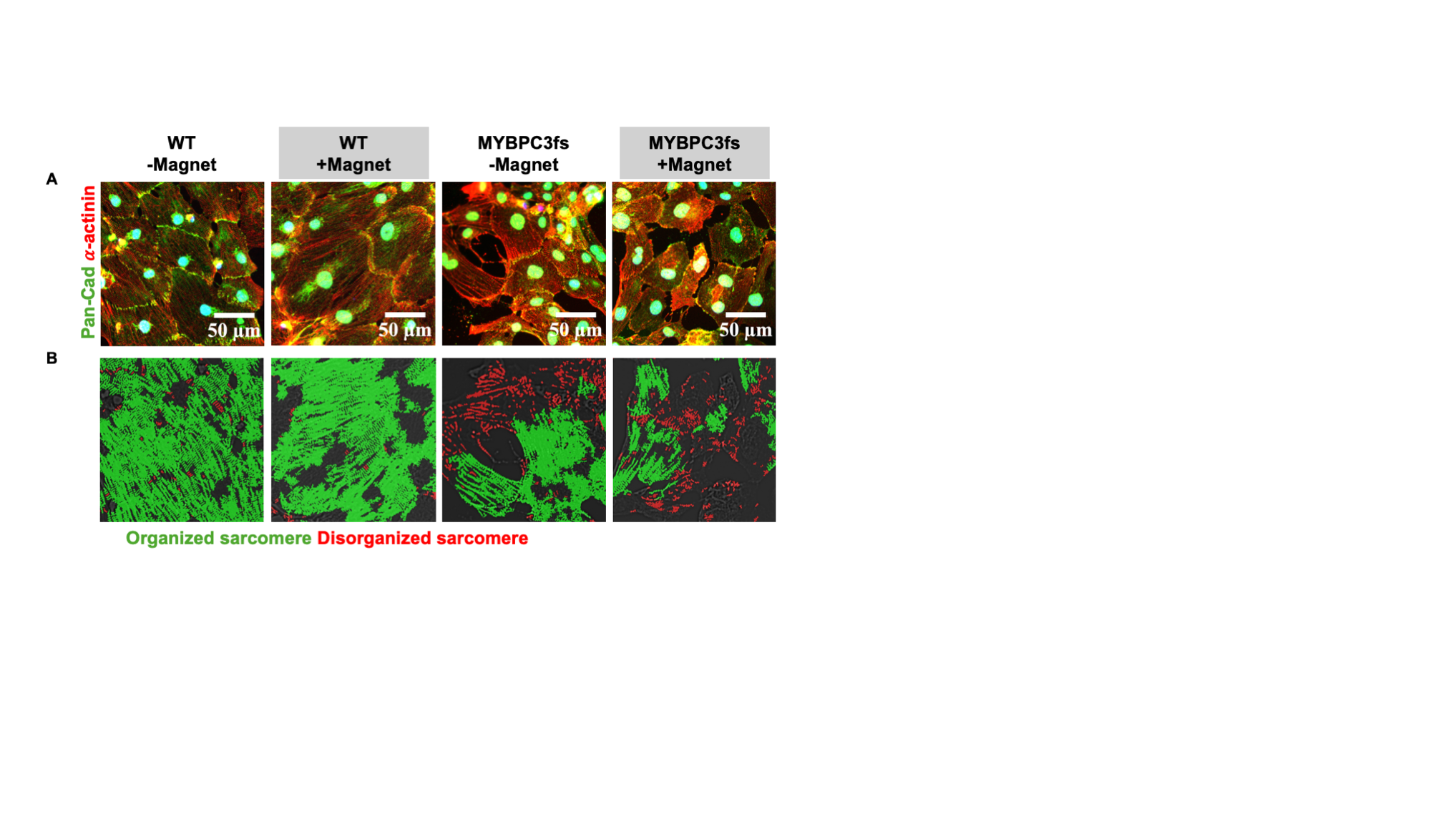

**Figure S9. SarcAsM-based sarcomere organization analysis reveals genotype- and mechanically-dependent disorganization in hiPSC-CMs.** **(A)** Representative immunofluorescence images of WT and *MYBPC3fs* hiPSC-CMs under unmagnetized (−Magnet) and magnetized (+Magnet) conditions, stained for Pan-Cadherin (green), 𝛼-actinin (red), and Hoechst-stained nuclei (blue). Scale bars: 50 𝜇m. **(B)** Corresponding SarcAsM analysis outputs overlaid on the 𝛼-actinin channel, pseudocolored to distinguish organized sarcomeres (green) from disorganized sarcomeres (red). WT unmagnetized cardiomyocytes show predominantly organized sarcomere structure, which is partially disrupted upon magnetization. *MYBPC3fs* cardiomyocytes show reduced baseline sarcomere organization compared to WT controls, which is further exacerbated by mechanical stimulation, with magnetized *MYBPC3fs* cells showing the greatest proportion of disorganized sarcomere area across all conditions.

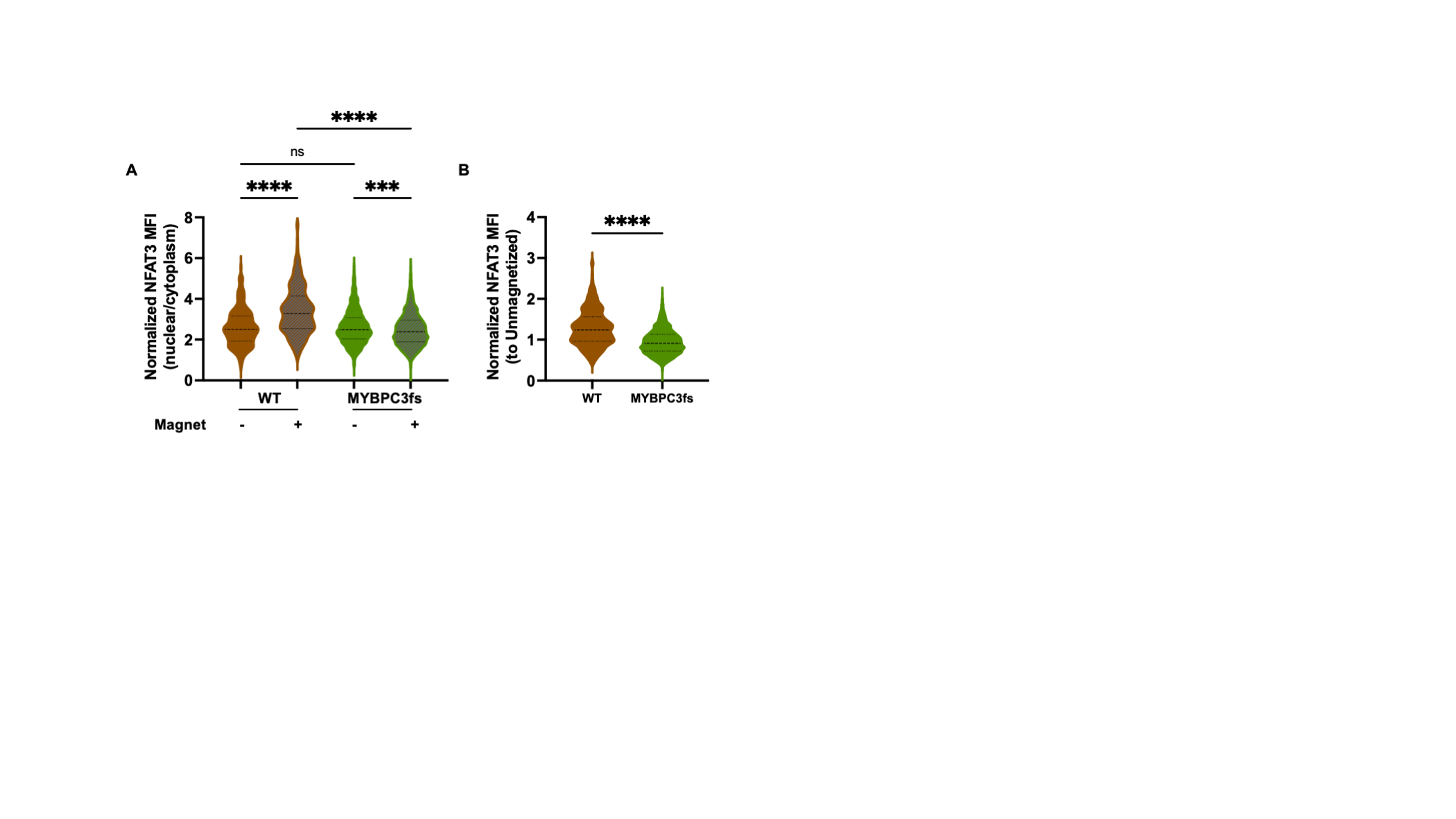

**Figure S10. Mechanical stimulation induces divergent NFAT3 subcellular redistribution in isogenic control and *MYBPC3fs* cardiomyocytes. (A)** NFAT3 nuclear-to-cytoplasmic MFI ratio across WT and *MYBPC3fs* hiPSC-CMs under unmagnetized (−) and magnetized (+) conditions. Magnetization significantly increased the nuclear-to-cytoplasmic ratio in WT and decreased in *MYBPC3fs* cardiomyocytes, with magnetized WT cells reaching significantly higher levels than magnetized *MYBPC3fs* cells. **(B)** Normalized (nuclear/cytoplasm) NFAT3 MFI relative to the respective unmagnetized condition, comparing WT and *MYBPC3fs* cardiomyocytes following magnetization. *MYBPC3fs* cardiomyocytes showed a significantly lower normalized (nuclear/cytoplasm) NFAT3 MFI compared to WT upon magnetization. Data are presented as violin plots, n represent individual cell; *P* values by one-way ANOVA, non-parametric, Kruskal-Wallis test with Dunn's multiple comparisons (A) and unpaired t-test, non-parametric, Mann-Whitney test (B); **** *p* < 0.0001, *** *p* < 0.001, ns = not significant.

1.	Härtter D, et al. “SarcAsM: AI-based multiscale analysis of sarcomere organization and contractility in cardiomyocytes”. *bioRxiv*. 2025:2025.04.29.650605.
